# The germline coordinates mitokine signaling

**DOI:** 10.1101/2023.08.21.554217

**Authors:** Koning Shen, Jenni Durieux, Cesar G. Mena, Brant M. Webster, C. Kimberly Tsui, Hanlin Zhang, Lawrence Joe, Kristen Berendzen, Andrew Dillin

## Abstract

The ability of mitochondria to coordinate stress responses across tissues is critical for health. In *C. elegans*, neurons experiencing mitochondrial stress elicit an inter-tissue signaling pathway through the release of mitokine signals, such as serotonin or the WNT ligand EGL-20, which activate the mitochondrial unfolded protein response (UPR^MT^) in the periphery to promote organismal health and lifespan. We find that germline mitochondria play a surprising role in neuron-to-peripheral UPR^MT^ signaling. Specifically, we find that germline mitochondria signal downstream of neuronal mitokines, like WNT and serotonin, and upstream of lipid metabolic pathways in the periphery to regulate UPR^MT^ activation. We also find that the germline tissue itself is essential in UPR^MT^ signaling. We propose that the germline has a central signaling role in coordinating mitochondrial stress responses across tissues, and germline mitochondria play a defining role in this coordination because of their inherent roles in germline integrity and inter-tissue signaling.

## Introduction

Multi-cellular organisms must coordinate the health and function of varied tissues across the organism in order to maintain overall health. This has led to the development of specialized tissues or organs that can communicate long-range signaling with other tissues to ensure organismal health. For instance, the central nervous system regulates the activity of other organs in the body through hormones or neuronal signaling (e.g. neurotransmitters and neuropeptides). Similarly, muscle or adipose fat can send metabolic or hormonal signals to other components of the body under conditions of hunger, overnutrition, or exercise.

These pathways of inter-tissue signaling become especially crucial during conditions of stress, such as pathogenic infection or cold exposure (Castillo-Armengol et al., 2019; Rao et al., 2017), or during unhealthy aging, where the aging-induced dysfunction in certain organs may contribute to loss of inter-tissue communication pathways and a broader loss of organismal health (Smith et al., 2020; Van Oosten-Hawle, 2023). For instance, over-accumulation of inflammation or senescent cells in certain tissues, such as muscle and the brain, can release blood-borne factors that affect the aging process (Bieri et al., 2023; Horowitz et al., 2020). In some cases, these secreted factors are protective, such as GDF11 which increases synaptic plasticity and neurogenesis (Bieri et al., 2023; Katsimpardi et al., 2014; Ozek et al., 2018). However, some secreted factors can also aggravate the aging process, such as the proinflammatory chemokines secreted by senescent cells (Birch and Gil, 2020), suggesting that inter-tissue signaling mechanisms must be tightly regulated to ensure organismal health.

Signals for inter-tissue communication can arise from a multitude of cellular sources, and mitochondria are a prominent source of these signals (Chandel, 2014; Mottis et al., 2019; Shen et al., 2022). This important signaling role for mitochondria may have arisen out of their endosymbiotic origin. When the prokaryotic mitochondrial precursor was engulfed by its now- eukaryotic host, the mitochondria had to develop comprehensive signaling pathways to communicate with its new host cell and nuclear genome. Fittingly, these mitochondrial signaling pathways are highly conserved and have been observed in species ranging from mammals through invertebrates. In particular, mitochondria experiencing dysfunction use inter-tissue signaling pathways to communicate their stress to other tissues and activate protective cellular programs, such as the mitochondrial unfolded protein response (UPR^MT^), a transcriptional program activated under conditions of mitochondrial stress to alleviate mitochondrial burden and promote overall cellular and organismal health (Houtkooper et al., 2013; Moehle et al., 2019; Shpilka and Haynes, 2018). In mice, low-level mitochondrial stress in POMC neurons induced by *Crif1* heterodeficiency signals to white adipose tissue to increase thermogenesis and activate the UPR^MT^ to protect against obesity and enhance metabolic turnover (Kang et al., 2021). In *Drosophila*, muscle tissue experiencing mild mitochondrial stress also activates the UPR^MT^ and induces expression of ImpL2 (an ortholog to the human insulin-like growth factor binding protein IGFBP7) that suppresses whole-organism insulin signaling, upregulates mitophagy, and promotes longevity and muscle function (Owusu-Ansah et al., 2013).

Much of the mechanistic work investigating inter-tissue mitochondrial signaling has been performed in *C. elegans* using a model of inter-tissue (or cell nonautonomous) communication of the UPR^MT^ between neurons and the intestine (Durieux et al., 2011). In this model, neurons experiencing mitochondrial stress, such as neuron-specific knockdown of mitochondrial genes (Durieux et al., 2011), expression of the aggregation-prone mutant PolyQ40 protein, or neuron-specific targeting of KillerRed to mitochondria (Berendzen et al., 2016), activate the UPR^MT^ in the intestine, a tissue in *C. elegans* completely peripheral and non-innervated by neurons **(Fig 1A)**.

**Figure 1:**
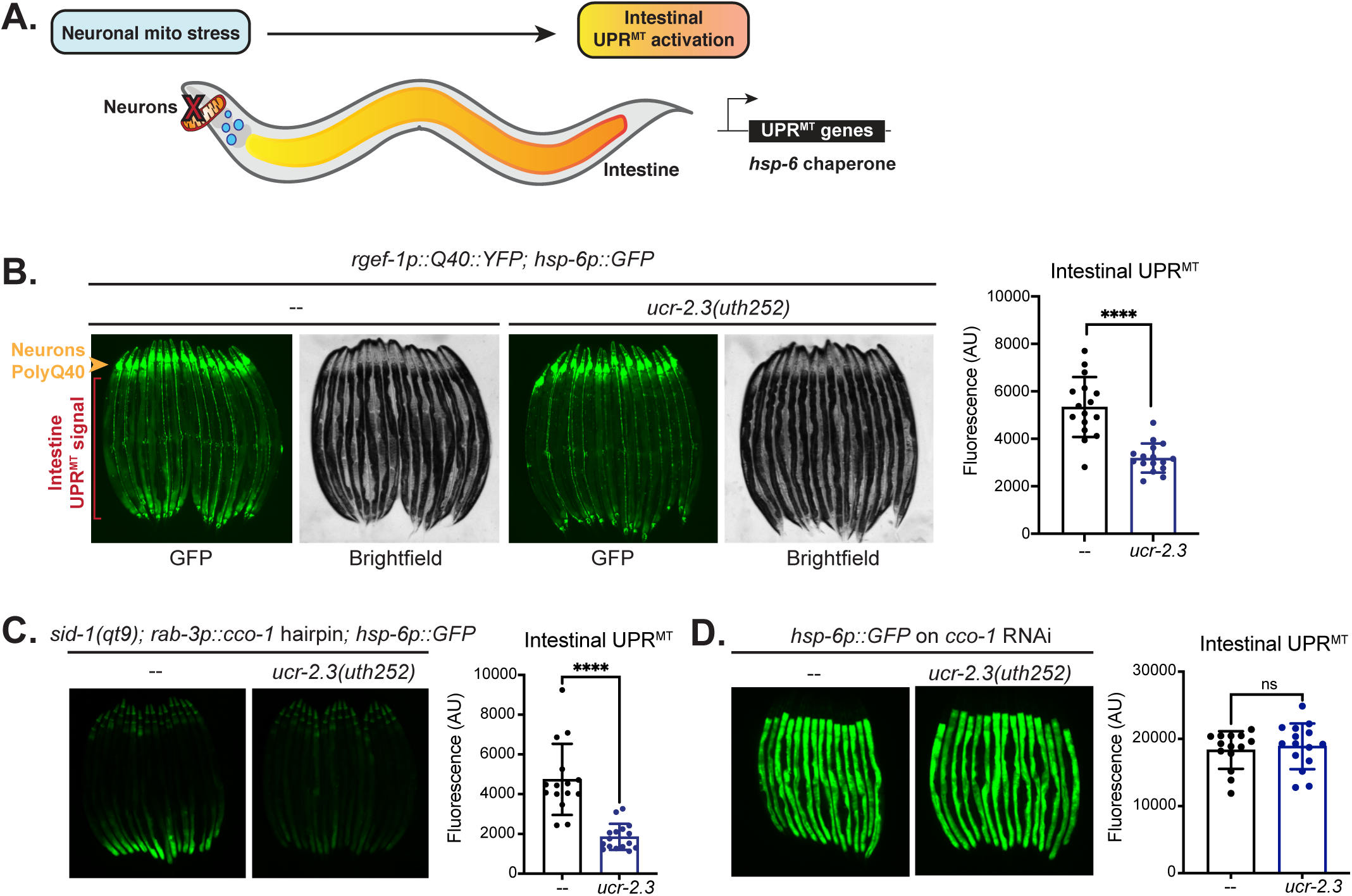
Mutagenesis screen reveals the requirement of *ucr-2.3* in neuron-to-intestine UPR^MT^ signaling. a. Schematic of cell nonautonomous UPR^MT^ signaling model in *C. elegans*. Through an inter- tissue signaling mechanism, neuron-specific mitochondrial stress activates the UPR^MT^ in the intestine of *C. elegans*. Neuronal mitochondrial stress can be induced by neuron- specific expression of a mutant, aggregation-prone PolyQ40 protein (*rgef-1p::Q40::YFP*) or neuron-specific knockdown mitochondrial genes like complex IV subunit *cco-1*. UPR^MT^ activation can be detected by use of a transcriptional reporter *hsp-6p::GFP*. See text for more details. b. Fluorescence imaging and quantification of *rgef-1p::Q40::YFP; hsp-6p::GFP* animals with and without loss of function mutation *ucr-2.3(uth252)*. The Q40::YFP signal in *C. elegans* neurons is denoted with an orange arrowhead. The UPR^MT^ transcriptional reporter signal (GFP expression driven by the *hsp-6* promoter) is denoted with the red brackets. Significance determined by Welch’s unpaired t-test, ****p<0.0001; n > 3. c. Fluorescence imaging and quantification of *sid-1(qt9); rab-3p::cco-1* hairpin*; hsp- 6p::GFP* animals with and without loss of function mutation *ucr-2.3(uth252)*. Significance determined by Welch’s unpaired t-test, ****p<0.0001; n > 3. d. Fluorescence imaging and quantification of *hsp-6p::GFP* animals with and without loss of function mutation *ucr-2.3(uth252)* fed *cco-1* RNAi . Significance determined by Welch’s unpaired t-test, p = 0.6372; n > 3.

Intestinal activation of UPR^MT^ by these neurons protected organismal health, as inhibition of UPR^MT^ activation decreased organismal fitness in the presence of neuronal mitochondrial stress (Berendzen et al., 2016).

Several neuronal factors mediating neuron-to-intestine UPR^MT^ signaling in *C. elegans* have been identified. Neuron-specific over-expression of the histone demethylases *jmjd-1.2* and *jmjd-3.1* induces intestinal UPR^MT^ even in the absence of mitochondrial stress and also regulates activation of UPR^MT^ in mice (Merkwirth et al., 2016). Neuronal release of neurotransmitters like serotonin or the neuropeptides FLP-1 and FLP-2 are also critical in mediating cell nonautonomous activation of the UPR^MT^ and other stress responses in the intestine (Berendzen et al., 2016; Jia and Sieburth, 2021; Shao et al., 2016). The WNT ligand, EGL-20, was identified as a bona fide “mitokine” as it is both necessary and sufficient for activation of the intestinal UPR^MT^ upon neuron-specific over-expression (Zhang et al., 2018). Neuronal release of the EGL-20 WNT ligand engages canonical WNT signaling pathway in the peripheral intestinal cells to activate the UPR^MT^ (Zhang et al., 2018). However, a dichotomy exists because the canonical developmental program regulated by WNT signaling is not subsequently induced. Therefore, how this WNT signaling, in the context of neuronal mitochondrial stress, could specifically induce the UPR^MT^ and not its canonical developmental programs in the intestine is unknown.

To address this question, we performed a genome-wide mutagenesis screen in *C. elegans* to identify molecular players that mediated UPR^MT^ cell nonautonomous signaling between neurons and the intestine. Through this screen, we identified mutations in the *ucr-2.3* gene that are essential in mediating cell nonautonomous UPR^MT^ signaling between neurons and the intestine. *ucr-2.3* is a mitochondrial electron transport chain (ETC) complex III gene, homologous to mammalian UQCRC2, and is expressed primarily in a few neurons and the germline of *C. elegans*. Surprisingly, we discovered that germline activity, not neuronal, of *ucr-2.3* mediates the cell nonautonomous UPR^MT^ signaling between the neurons and intestine. In addition, we find that the germline plays a general role in mediating cell nonautonomous UPR^MT^ signaling, as loss of the germline through either genetic or pharmacological treatments abolishes neuron-to-intestine UPR^MT^ activation, but not other neuronal derived stress signals. These data suggest that integrity of the germline tissue, mediated through germline UCR-2.3 mitochondrial activity, is integral for neuron-to-intestinal signaling of the UPR^MT^. Finally, we find that germline integrity may regulate intestinal UPR^MT^ activation through altering lipid metabolism and transport pathways between reproductive and intestine tissues. Overall, our work showcases a signaling role for the germline, once thought of as a tissue insulated from the soma, in directly communicating with somatic tissues for stress signaling. Additionally, our results provide broader implications for the importance of germline mitochondrial quality and its physiological interactions with neuronal signaling and intestinal lipid metabolism, and the consequences of these interactions on organismal stress signaling and fertility.

## Results

### Mutagenesis screen reveals the requirement of *ucr-2.3* in neuronal UPR^MT^ signaling

To identify novel regulators of inter-tissue (or cell nonautonomous) mitochondrial signaling in *C. elegans*, we conducted a genome-wide mutagenesis screen using a genetic model where neurons expressing an aggregation-prone mutant PolyQ40 protein tagged with YFP (*rgef- 1p::Q40::*YFP) activates the mitochondrial unfolded protein response (UPR^MT^) in the intestine (Berendzen et al., 2016) **(Fig 1A)**. Activation of the UPR^MT^ in the intestine can be assessed by a transcriptional reporter for expression of *hsp-6*, the mitochondrial Hsp70 chaperone induced by the UPR^MT^ (Shpilka and Haynes, 2018) **(Fig 1A)**. After identifying mutants with suppressed intestinal UPR^MT^ but intact neuronal PolyQ40 signal, we used SNP mapping and whole genome sequencing approaches (Doitsidou et al., 2010) to identify one causative mutation that suppressed cell nonautonomous UPR^MT^ signaling as a missense mutation in the gene *ucr-2.3* **(Supp Fig 1A, F)**. We independently verified the necessity of *ucr-2.3* in mediating cell nonautonomous UPR^MT^ signaling by creating several alleles of early stop codon, nonsense-mediated decay (NMD) mutations in *ucr-2.3* through CRISPR/Cas9 genome editing and observed that they all suppressed PolyQ40-mediated cell nonautonomous UPR^MT^ **(Fig 1B; Supp Fig 1B, F)**. Importantly, the NMD early stop codon mutation in the *ucr-2.3(uth252)* allele leads to almost complete loss of gene expression, thus effectively creating a null allele **(Supp Fig 1D, F)**. We also tested a *ucr-2.3* mutant containing a large deletion spanning exons 3-4 resulting in an early termination codon and found that it also suppressed cell nonautonomous UPR^MT^ signaling **(Supp Fig 1C, F).** We also tested whether *ucr-2.3* was involved in other models of cell nonautonomous UPR^MT^ signaling, such as neuron-specific knockdown of the mitochondrial complex IV subunit *cco-1* (*rab-3p::cco-1* hairpin in the *sid-1(qt9)* genetic background), which also activates UPR^MT^ in the intestine (Durieux et al., 2011). We find that a null mutation in *ucr-2.3* also suppresses this model of cell nonautonomous UPR^MT^ signaling **(Fig 1C)**. Therefore, we conclude that *ucr-2.3* is as a novel regulator of neuron-to-intestine UPR^MT^ signaling induced by multiple forms of neuronal mitochondrial stress.

Next, we tested whether *ucr-2.3* was required for cell autonomous induction of the UPR^MT^. We tested this by feeding *cco-1* RNAi targeting a mitochondrial complex IV subunit to wildtype and *ucr-2.3* mutant strains to ask whether loss of *ucr-2.3* impairs cell-autonomous activation of the UPR^MT^. We find that *ucr-2.3* loss of function mutants still activate, not inhibit, the UPR^MT^ cell autonomously, with and without neuronal PolyQ40 expression **(Fig 1D, Supp Fig 1E)**. These data indicate that *ucr-2.3* is not involved in general machinery of UPR^MT^ activation but rather has a specific role in cell nonautonomous signaling of mitochondrial stress. We also find that loss of *ucr-2.3* non-autonomous UPR^MT^ signaling is not a temporary delay. Rather, the suppression of non-autonomous UPR^MT^ signaling by the *ucr-2.3* mutation increases over aging **(Supp Fig 1G)**. Finally, we tested the specificity of *ucr-2.3* loss in nonautonomous stress signaling and find that *ucr-2.3* loss of function does not suppress cell nonautonomous signaling of the UPR^ER^ or the cytosolic heat stress response (HSR) between neurons and the intestine **(Supp Fig 2A-B)**. Conversely, there is a slight up-regulation of both stress responses upon *ucr-2.3* loss of function. Overall, our data indicate that *ucr-2.3* is a critical and specific gene required for mediating cell nonautonomous UPR^MT^ signaling between neurons and the intestine in *C. elegans*.

### UCR-2.3 is a mitochondrial protein with a novel role in mitochondrial stress signaling

The *ucr-2.3* gene encodes UCR-2.3 in *C. elegans*, which is reported to be a mitochondrial protein homologous to mammalian UQCRC2, a core subunit of respiratory complex III of the electron transport chain (Iwata et al., 1998; Nomura et al., 2006). That the loss of *ucr-2.3* results in a loss of UPR^MT^ neuronal signaling was perplexing because the knockdown of many mitochondrial proteins, especially those involved in the electron transport chain, activate, not suppress, the UPR^MT^ (Durieux et al., 2011; Houtkooper et al., 2013; Nargund et al., 2012). Indeed, we observed that even in “basal” conditions, i.e. no additional source of stress, the *ucr-2.3* null mutation still does not activate UPR^MT^ **(Supp Fig 2C)**, suggesting that UCR-2.3 in mitochondria may have a non-canonical role in regulating UPR^MT^ signaling.

We experimentally confirmed that the UCR-2.3 protein is a mitochondrial protein using three orthogonal approaches. First, we performed a sub-cellular fractionation in *C. elegans* expressing *ucr-2.3* with an HA epitope endogenously inserted at the *ucr-2.3* genomic locus using CRISPR/Cas9 editing **(Fig 2A*i*)**. Compared to the cytosolic fraction, we detected enrichment of UCR-2.3 in the mitochondrial fraction similar to that of NDUFS3, a subunit of mitochondrial electron transport complex I, and as opposed to α-tubulin, which was detected predominantly in the cytosolic fraction **(Fig 2A*ii*)**. Second, using human RPE-1 cells, we over-expressed the *C. elegans* UCR-2.3 protein tagged with mNeonGreen alongside an outer mitochondrial membrane protein OMP25 tagged with BFP **(Fig 2B)**. We find that UCR-2.3 and OMP-25 thoroughly co- localize (mean 71.8%, SD = 18.2%), indicating the UCR-2.3 is highly localized to mitochondria **(Fig 2B*i*)**. Indeed, upon closer examination, we observed that UCR-2.3 primarily localizes within the OMP25 signal, suggesting that UCR-2.3 is located within the mitochondrial matrix, consistent with its purported role as a matrix-facing core subunit of complex III like UQCRC2 **(Fig 2B*ii*)** (Iwata et al., 1998). Third, we observed that UCR-2.3 functionally acted as a mitochondrial protein. Using the null mutation in *ucr-2.3* in *C. elegans*, we observed that loss of *ucr-2.3* decreased mitochondrial respiration **(Fig 2C)**. Thus, we conclude that *ucr-2.3* encodes a bona fide mitochondrial protein in *C. elegans*, which has a similar sub-cellular localization and function to its mammalian ortholog UQCRC2.

**Figure 2:**
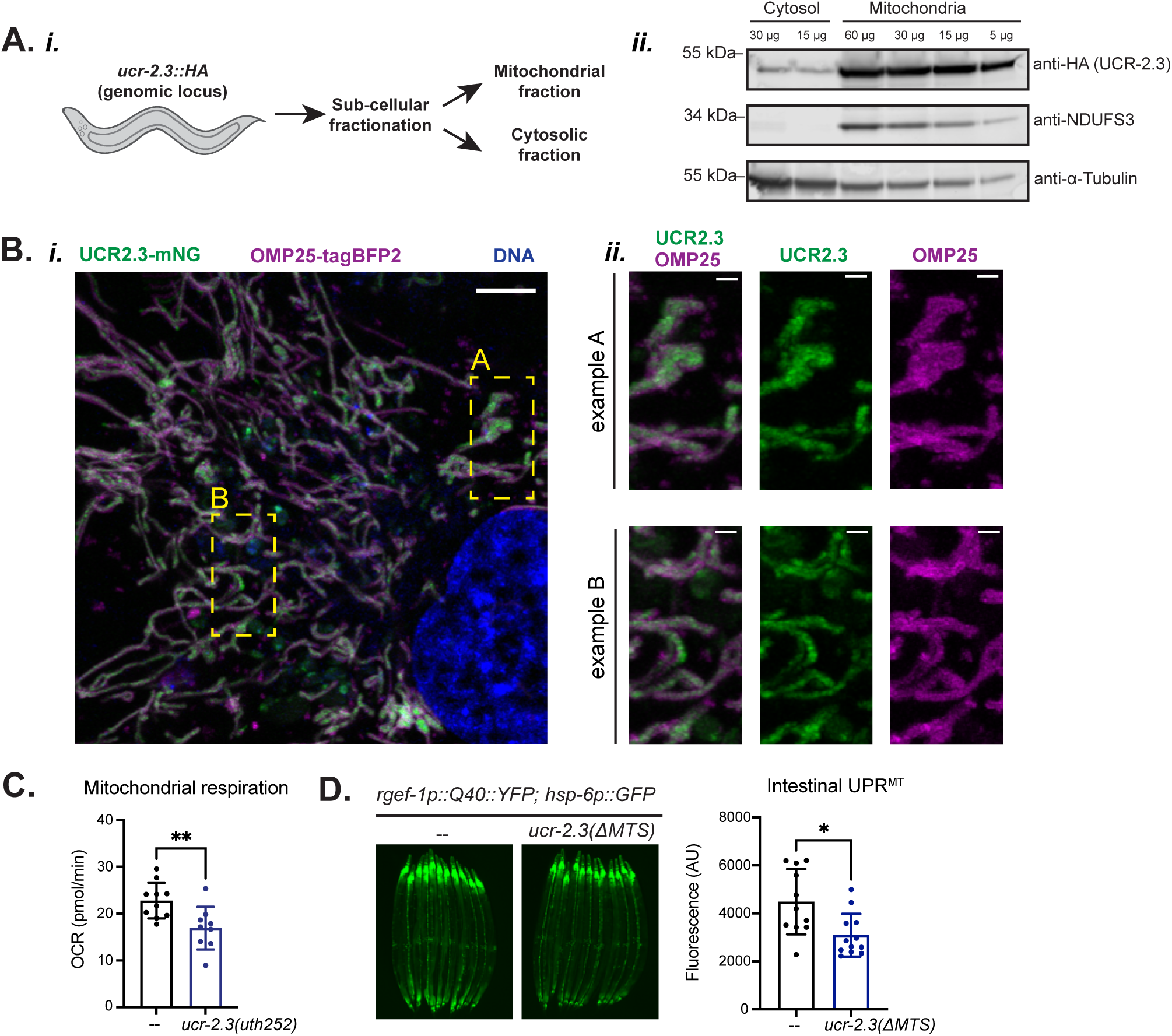
UCR-2.3 is a mitochondrial protein in *C. elegans* with a novel role in cell nonautonomous mitochondrial stress signaling. a. *(i)* Schematic of sub-cellular fractionation of a *C. elegans* strain with a HA-tag inserted at the C-terminus of the *ucr-2.3* genomic locus using CRISPR/Cas9 genomic editing. Cytosolic and mitochondrial fractions were run on a western blot probed with an HA antibody (to detect UCR-2.3 protein), a NDUFS3 antibody (to detect an established mitochondrial ETC protein), and an α-tubulin antibody (to detect an established cytosolic protein). b. *(i)* Live-cell imaging showing sub-cellular localization of *C. elegans* UCR-2.3 in human RPE1 hTert cells transduced with *C. elegans* UCR2.3 tagged with mNeonGreen (UCR- 2.3-mNG) and OMP25 tagged with BFP (OMP25-tagBFP2). Scale bar is 5 μm. *(ii)* Close up of two example regions shown in *(i)*. Scale bar is 1 μm. Data shown is representative of at least two independent imaging experiments with 5-10 images collected per cell line. c. Measurement of mitochondrial respiration or OCR (oxygen consumption rate) between wildtype and the *ucr-2.3(uth252)* mutant in the *rgef-1p::Q40::YFP; hsp-6p::GFP* genetic background. Significance determined by Welch’s unpaired t-test, ** p = 0.0079; n > 4. d. Fluorescence imaging comparison of intestinal UPR^MT^ between wildtype and a *ucr-2.3* mutant with the predicted mitochondrial targeting sequence (MTS) deleted *ucr-2.3(11MTS)*, all in the *rgef-1p::Q40::YFP; hsp-6p::GFP* genetic background. Significance determined by Welch’s unpaired t-test, * p = 0.0103; n = 3.

Next, we tested whether UCR-2.3 mitochondrial localization was essential for its role in mediating cell nonautonomous UPR^MT^ signaling. Using CRISPR/Cas9 genomic editing, we excised the predicted mitochondrial targeting sequence (MTS) of UCR-2.3 predicted by MitoFates (Fukasawa et al., 2015) from UCR-2.3 and found that this ΔMTS mutation *ucr-2.3(ΔMTS)* suppressed intestinal UPR^MT^ signaling **(Fig 2D)** but had no effect on *ucr-2.3* expression **(Supp Fig 2D)**. Overall, we conclude that *ucr-2.3* encodes a mitochondrial localized protein whose mitochondrial activity is important for regulating neuron-to-intestine cell nonautonomous UPR^MT^ signaling in a manner atypical of canonical mitochondrial electron transport chain proteins.

### *ucr-2.3* has a unique role within the UCR-2 gene family in *C. elegans*

*ucr-2.3* belongs to a family of paralogs in the UCR-2 family in *C. elegans* that are all homologous to mammalian UQCRC2: *ucr-2.1, ucr-2.2,* and *ucr-2.3* **(Fig 3A)** (Howe et al., 2016). To investigate whether the other paralogs of the UCR-2 family were also involved in cell nonautonomous UPR^MT^ signaling, we made similar early stop codon NMD mutations in *ucr-2.1* and *ucr-2.2* using CRISPR/Cas9 editing **(Supp Fig 3E*i-ii*)**. Surprisingly, we found that only genetic loss of *ucr-2.3* suppressed cell nonautonomous UPR^MT^ signaling **(Fig 3B)**. Genetic loss of *ucr-2.1* or *ucr-2.2* slightly activated, not suppressed, UPR^MT^, more in alignment with the expectation of a canonical mitochondrial gene knockdown **(Fig 3B)**. This suggested that *ucr-2.3* has a unique role among the UCR-2 family genes, whereas *ucr-2.1* and *ucr-2.2* may be more similar and typical of electron transport chain subunits.

**Figure 3:**
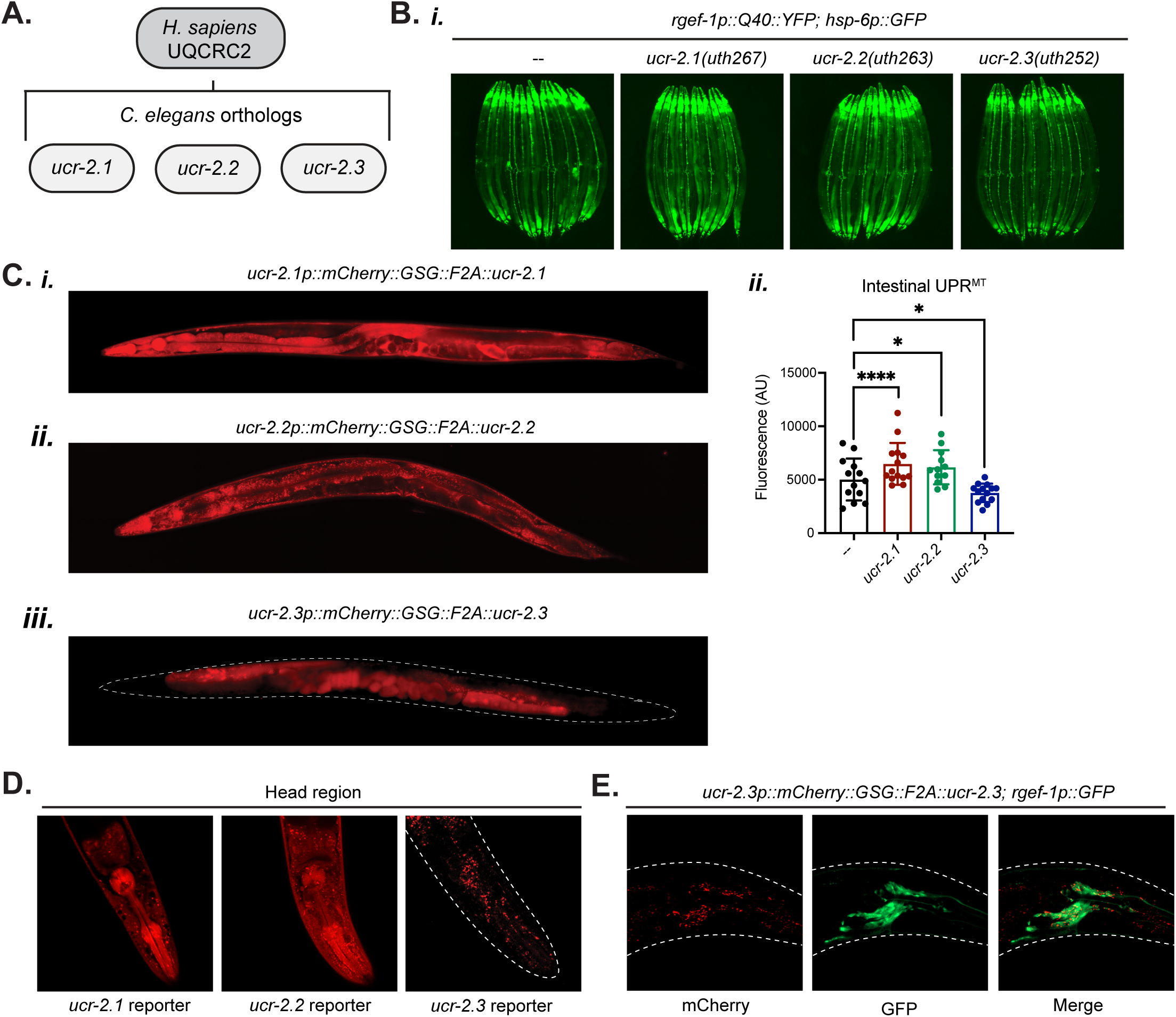
*ucr-2.3* is part of the UCR-2 family of gene paralogs that differ in their tissue expression and role in cell nonautonomous UPR^MT^ signaling. a. Human UQCRC2 is homologous to a family of UCR-2 genes in *C. elegans* including *ucr- 2.1*, *ucr-2.2*, and *ucr-2.3*. b. *(i)* Fluorescence imaging of comparing the loss of function mutations in the UCR-2 family on cell nonautonomous UPR^MT^ signaling. All mutations showed are nonsense-mediated decay mutations generated via CRISPR/Cas9 editing. Quantification (*ii*). Significance determined by Welch’s unpaired t-test. *p < 0.05, ****p < 0.0001; n = 3. c. Fluorescence imaging comparison of tissue expression patterns across the UCR-2 family genes using an endogenous transcriptional reporter for each gene: *ucr-2.1* (*i*), *ucr-2.2* (*ii*), and *ucr-2.3* (*iii*). To generate each transcriptional reporter, a mCherry::GSG linker::F2A sequence was inserted between the gene promoter and coding region at the genomic locus using CRISPR/Cas9 editing. d. Close up head images of the endogenous transcriptional reporter as described in (C). e. Close up head imaging of *ucr-2.3* endogenous transcriptional reporter crossed to a *rgef- 1p::GFP* reporter strain, expressing GFP in neurons.

Next, we next sought to explore the functional and genetic relationships among the UCR- 2 family genes. First, we observed that *ucr-2.1* and *ucr-2.2* were functionally redundant with each other with respect to their role in the mitochondrial electron transport chain, but neither were redundant with *ucr-2.3.* While knockdown of *ucr-2.3* reduced mitochondrial respiration **(Fig 2C)**, only the double knockdown of *ucr-2.1* and *ucr-2.2* could reduce mitochondrial respiration **(Supp Fig 3A)**. Individual knockdown of either *ucr-2.1* or *ucr-2.2* had no effect on respiration, suggesting that these two genes are functionally redundant with each other **(Supp Fig 3A)**. In addition, double knockdown of *ucr-2.1* and *ucr-2.2* strongly activated UPR^MT^, while the single knockdowns had no effect **(Supp Fig 3B)**. As this strong activation is consistent with the expectation of a canonical mitochondrial ETC gene knockdown (Durieux et al., 2011; Houtkooper et al., 2013), this provides further evidence that *ucr-2.1* and *ucr-2.2* are functionally redundant with each other for a role in the mitochondrial electron transport chain, likely together recapitulating the role of the mammalian UQCRC2 complex III subunit. In contrast, we observed that double knockdown of either *ucr-2.1* and *ucr-2.3* or *ucr-2.2* and *ucr-2.3* could not activate UPR^MT^; only conditions in which both *ucr-2.1* and *ucr-2.2* are knocked down could UPR^MT^ be activated **(Supp Fig 3C)**. This finding further suggests that neither *ucr-2.1* nor *ucr-2.2* are functionally redundant with *ucr-2.3*.

In addition, we analyzed how knockdown of these UCR-2 family genes affected each other transcriptionally. We found that while knockdown of *ucr-2.3* increased expression of *ucr-2.1* and *ucr-2.2* **(Supp Fig 3D)**, knockdown of either *ucr-2.1* or *ucr-2.2* individually or together had no effect on *ucr-2.3* expression **(Supp Fig 3E)**. One interpretation of this result is that loss of *ucr-2.3* function requires compensation from both *ucr-2.1* and *ucr-2.2* equally, while loss of either *ucr-2.1* or *ucr-2.2* does not require *ucr-2.3.* Altogether, these data suggest that *ucr-2.1* and *ucr-2.2* are functionally redundant with each other while *ucr-2.3* may have an independent role.

Most importantly, we found that the UCR-2 family genes differ in their tissue expression patterns **(Fig 3C-D)**. Using genomic, tissue-specific, and single cell transcriptomic databases, we found that while *ucr-2.1* and *ucr-2.2* expression is mostly ubiquitous through the animal, *ucr-2.3* expression is enriched in the germline and neuronal cells (Cao et al., 2017; Howe et al., 2016; Kaletsky et al., 2018) **(Supp Fig 3F)**. We confirmed these reports by generating an endogenous transcriptional expression reporter system for each of the UCR-2 genes by inserting an mCherry::F2A construct between the promoters of *ucr-2.1*, *ucr-2.2*, and *ucr-2.3* and the respective CDS at the endogenous genomic loci using CRISPR/Cas9 editing **(Fig 3C)**. The tissue expression patterns confirm that *ucr-2.1* and *ucr-2.2* are mostly ubiquitously expressed throughout the somatic tissues of the animal, though largely excluded from the germline **(Fig 3C*i-ii*, 3D)**. In contrast, *ucr-2.3* expression was exclusively localized to the germline as well as a few cells in the head of the animal **(Fig 3C*iii*, 3D)**. To determine which of these head cells expressed *ucr-2.3*, we crossed the *ucr-2.3* expression reporter strain to an *rgef-1p::GFP* construct driving GFP expression in neurons and found that the pattern of *ucr-2.3* expression mostly overlapped with the GFP signal **(Fig 3E)**, suggesting that the cells expressing *ucr-2.3* are indeed neurons. In summary, our data indicate that *ucr-2.3* differs starkly in its tissue expression from *ucr-2.1* and *ucr-2.2*, which may explain the unique role of *ucr-2.3* in mediating cell nonautonomous UPR^MT^ signaling.

### The germline role of *ucr-2.3* is critical for mediating nonautonomous UPR^MT^

We were intrigued by the tissue-specific expression patterns of *ucr-2.3* and tested whether this tissue-specific nature conferred the ability of *ucr-2.3* to mediate cell nonautonomous UPR^MT^ signaling. To do this, we sought to determine the site of action for *ucr-2.3* for its role in cell nonautonomous UPR^MT^ signaling. We designed a tissue-specific rescue experiment using the single copy mosSCI system (Frøkjær-Jensen et al., 2008) **(Fig 4A)**. Briefly, we knocked in an integrated wildtype copy of *ucr-2.3* driven by a tissue-specific promoter into a mutant strain of *ucr-2.3*. As we observed that *ucr-2.3* is expressed in neuronal cells and the germline **(Fig 3C-D)**, we tested whether either neuronal or germline-specific expression of wildtype *ucr-2.3* could rescue cell nonautonomous UPR^MT^ signaling in the *ucr-2.3* mutant, using the neuron-specific promoter *rgef-1* and the germline-specific promoter *pie-1* **(Fig 4A)**. We found that expression of a wildtype copy of *ucr-2.3* driven by the germline-specific promoter *pie-1* could completely rescue the cell nonautonomous UPR^MT^ signal suppressed by loss of *ucr-2.3*, whereas *rgef-1p* neuronal-driven expression of *ucr-2.3* had no appreciable effect on cell nonautonomous UPR^MT^ signal **(Fig 4B)**. In addition, one of our initial observations of the *ucr-2.3* mutants was their reduced brood size (number of progeny) **(Fig 4C)**. We found that germline rescue of *ucr-2.3* could also completely restore the brood size phenotype we observed with *ucr-2.3* mutants, whereas neuronal rescue of *ucr-2.3* had no effect on the brood size **(Fig 4C)**.

**Figure 4:**
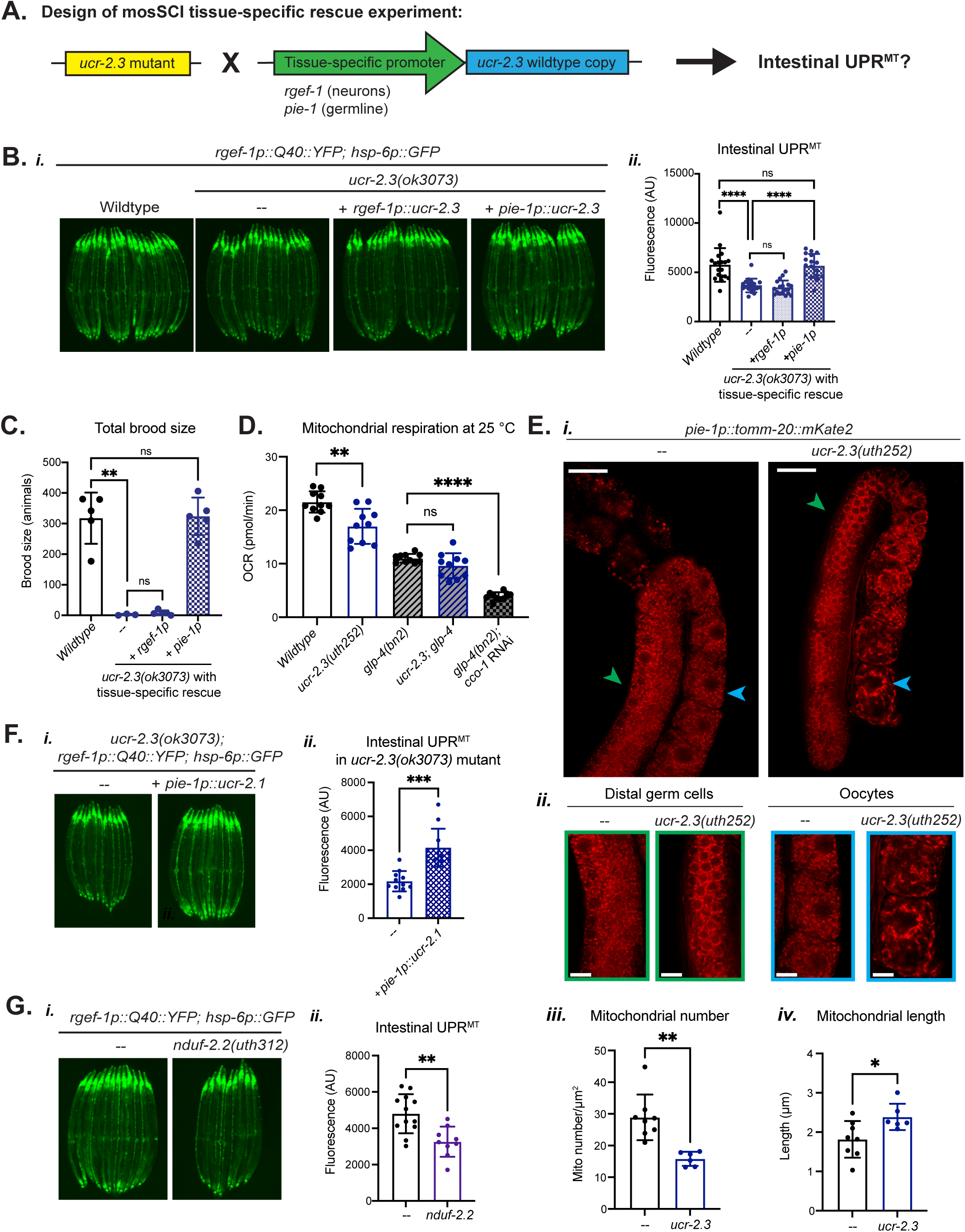
*ucr-2.3* acts in the germline to mediate cell nonautonomous UPR^MT^ signaling and germline mitochondrial integrity. a. Schematic of single-copy (mosSCI) tissue-specific rescue experiment. *ucr-2.3(ok3073)* mutant was crossed to strains containing an integrated, single-copy (mosSCI) of wildtype *ucr-2.3* driven by either a neuron-specific (*rgef-1p*) or germline-specific (*pie-1p*) promoter, all in the *rgef-1p::Q40::YFP; hsp-6p::GFP* genetic background. b. Fluorescence imaging comparison of tissue-specific mosSCI rescue experiment in the *rgef- 1p::Q40::YFP; hsp-6p::GFP* genetic background for the *ucr-2.3(ok3073)* mutation with tissue-specific rescue of wild-type *ucr-2.3* expressed in either in neurons (+ *rgef-1p*) or the germline (+ *pie-1p*) *(i)*. Significance determined by Welch’s unpaired t-test, ****p< 0.0001. All non-significant p values > 0.3924. Quantification *(ii)*. n = 3. c. Measurement of total brood size among wildtype, *ucr-2.3(ok3073)* loss of function mutant, and tissue-specific rescue with wild-type *ucr-2.3* expressed either in neurons (*rgef-1p::ucr- 2.3*) or the germline (*pie-1p::ucr-2.3*), all in the *rgef-1p::Q40::YFP; hsp-6p::GFP* genetic background. Significance determined by Welch’s unpaired t-test, **p=0.0011. All non- significant p values > 0.1507. n = 3. d. Measurement of mitochondrial respiration (oxygen consumption rate) for wildtype and the *ucr-2.3(uth252)* mutant with and without an intact germline, all in the *rgef-1p::Q40::YFP; hsp-6p::GFP* genetic background. The germline was genetically ablated via the *glp-4(bn2)* temperature-sensitive mutant strain and conducting the entire experiment at the restrictive temperature 25°C. All strains were fed control HT115 RNAi bacteria, except for the *glp- 4(bn2)* mutant animals fed *cco-1* RNAi. Significance determined by Welch’s unpaired t- test, **p = 0.0019, ****p < 0.0001, non-significant p = 0.1051. n = 3. e. Fluorescence widefield imaging of *pie-1p::tomm-20::mKate2* germline mitochondrial reporter strain with and without the *ucr-2.3(uth252)* mutation *(i)*. Scale bar = 25 μm. Distal germ cells (green arrowhead) and oocytes (blue arrowhead) are shown in zoomed in images *(ii)*. Scale bar = 10 μm. Images shown are representative of at least 5-10 animals imaged per strain in at least three independent experiments each. Quantification of average mitochondrial number *(iii)* and mitochondrial length *(iv)* between the wildtype and *ucr- 2.3(uth252)* mutant strain. Significance determined by Welch’s unpaired t-test, **p = 0.0010, *p = 0.0205; n > 2. f. Fluorescence imaging comparison of the *ucr-2.3(ok3073); rgef-1p::Q40::YFP; hsp- 6p::GFP* with and without the mosSCI rescue of germline-expressed wildtype *ucr-2.1* (*pie-1p::ucr-2.1*). Significance determined by Welch’s unpaired t-test, ***p=0.001; n = 2. g. Fluorescence imaging comparison *(i)* and quantification *(ii)* of *rgef-1p::Q40::YFP;hsp- 6p::GFP* with and without the *nduf-2.2(uth312)* null mutation. *nduf-2.2* has enriched expression to germline and neuronal tissues compared to its more ubiquitously expressed counterpart *gas-1*. Significance determined by Welch’s unpaired t-test, **p = 0.0015, n > 3.

We also used the FLP/FRT recombinase system to test how a neuron-specific knockout of *ucr-2.3* affected nonautonomous UPR^MT^ signaling (Muñoz-Jiménez et al., 2017). We were unable to use this system to test germline-specific knockout because germline-specific FLP/FRT recombination becomes pervasive in all tissues in the following generation (Macías-León and Askjaer, 2018). To design a neuron-specific FLP/FRT recombinase knockout system, we used the *rgef-1p* neuronal promoter to drive FLP recombinase expression and inserted FRT sites in the introns of *ucr-2.3* at the genomic locus. In this way, FLP recombination of *ucr-2.3* induced a large deletion in the *ucr-2.3* genomic locus only in neuronal cells, which we confirmed by PCR genotyping **(Supp Fig 4A*iii*)**. While *ucr-2.3* containing FRT sites had slightly reduced cell nonautonomous UPR^MT^ signal, we find that FLP induced recombination of *ucr-2.3* did not enhance reduction of the cell nonautonomous UPR^MT^ signal and did not phenocopy the loss of function *ucr-2.3* genetic mutant **(Supp Fig 4A*i-ii*)**. Overall, these data identify that *ucr-2.3* mediates cell nonautonomous UPR^MT^ signaling through its activity in the germline tissue.

Having previously discovered a role for *ucr-2.3* in mitochondrial respiration **(Fig 2C)**, we next asked how much of the *ucr-2.3* mitochondrial activity is confined to the germline tissue. We tested this by using a *C. elegans* strain with a temperature sensitive mutation in the *glp-4* gene, which arrests germ cell proliferation at the restrictive temperature, 25 °C, to create germline deficient adult animals (Beanan and Strome, 1992). We find that at the restrictive temperature 25 °C, *ucr-2.*3 loss of function still decreases mitochondrial respiration **(Fig 4D)**. Thus, germline deficient animals also have decreased mitochondrial respiration, likely due to the loss of the high population of mitochondria in the germline **(Fig 4D)** (Tsang and Lemire, 2002). However, additional loss of *ucr-2.3* function in the germline deficient animals had no impact on mitochondrial oxygen consumption **(Fig 4D)**. In contrast, feeding germline deficient animals with a RNAi targeting the *cco-1* complex IV ETC subunit could further decrease mitochondrial respiration **(Fig 4D)**. This indicates that the vast majority of *ucr-2.3* contribution to mitochondrial respiration is in the germline.

In addition, we find that loss of *ucr-2.3* greatly changes morphology and content of germline mitochondria. Using a germline-specific mitochondrial morphology reporter strain (Ahier et al., 2018), we observe that wildtype animals have a dense network of mitochondria throughout the germline tissue **(Fig 4E)**. However, the *ucr-2.3* mutation results in a more punctate and sparser mitochondrial network in the distal germ cells and especially oocytes **(Fig 4E)**. In addition, we find that while the average mitochondrial number (or mitochondrial density) decreases in the *ucr-2.3* mutant **(Fig 4E*iii*)**, average mitochondrial length increases **(Fig 4E*iii*)**. This result aligns with the idea that damaged mitochondria fuse together as compensation to increase overall oxidative capacity (Youle and Van Der Bliek, 2012), as *ucr-2.3* loss of function animals have compromised ETC activity **(Fig 2C**, **Fig 4D)**. Therefore, loss of *ucr-2.3* directly affects the quality of germline mitochondria, and that loss of this germline mitochondrial quality may compromise cell nonautonomous UPR^MT^ signaling.

Finally, we were intrigued by the germline tissue-specificity of *ucr-2.3* expression, as compared to its paralogs *ucr-2.1* and *ucr-2.2* or other mitochondrial ETC proteins. We asked whether nonautonomous UPR^MT^ signaling specifically required *ucr-2.3* activity or germline mitochondrial quality more generally. To answer this, we first tested whether germline-specific expression of the other *ucr-2* family genes, such as *ucr-2.1*, could rescue the *ucr-2.3* mutant suppression of nonautonomous UPR^MT^. Using the same single copy mosSCI system, we knocked in an integrated wildtype copy of *ucr-2.1* driven by the *pie-1* germline-specific promoter into a mutant strain of *ucr-2.3* **(Fig 4F)**. Surprisingly, we observed that germline-specific expression of *ucr-2.1* could also rescue nonautonomous UPR^MT^ signaling in the *ucr-2.3* loss of function mutant. Given the high sequence similarity between these two genes and their proposed similar function in complex III of the electron transport chain as the UQCRC2 *C. elegans* homolog (Nomura et al., 2006), this result suggests that general mitochondrial quality in the germline is essential for nonautonomous UPR^MT^ signaling.

As an additional measure of germline mitochondrial quality, we asked whether knockdown of mitochondrial genes that shared comparable tissue-specific paralogs as *ucr-2.3* could recapitulate the UPR^MT^ suppression we observed with *ucr-2.3* mutants. Through searching the same genomic and transcriptomic databases, we identified other mitochondrial electron transport chain subunits that had similarly predicted patterns of gene expression in which one paralog was ubiquitously expressed and another paralog was found restricted to neuronal and/or germline expression. One such pairing we found was the complex I subunit gene *gas-1/nduf-2.1* (ubiquitously expressed) and its related paralog *nduf-2.2* (neuron/germline expressed) **(Supp Fig 4B)** (Cao et al., 2017; Howe et al., 2016; Kaletsky et al., 2018). Similar to *ucr-2.3*, we used CRISPR/Cas9 genetic engineering to create early stop codon NMD mutations in the *nduf-2.2* genomic locus in the *rgef-1p::Q40::YFP; hsp-6p::GFP* genetic background. We observed that early stop NMD mutation in *nduf-2.2* also suppressed cell nonautonomous UPR^MT^ signaling but had no affect on autonomous UPR^MT^ signaling **(Fig 4G, Supp Fig 4C)**, similar to what we had observed for *ucr-2.3* mutations **(Fig 1B, Supp Fig 1E)**. Therefore, general germline mitochondrial dysfunction inhibits neuronal derived cell nonautonomous UPR^MT^ signaling.

### The germline, and germline mitochondria, is required for cell nonautonomous UPR^MT^ signaling

Given the essential role of *ucr-2.3* and mitochondria in the germline, we next asked if simply general germline function was also essential for mediating cell nonautonomous UPR^MT^ signaling. We tested this using genetic and pharmacological methods to deplete the germline in *C. elegans*. For genetics, we generated germline deficient animals using two temperature sensitive mutations *glp-4(bn2)* and *glp-1(e2141ts)*, which both restrict germ cell proliferation at the restrictive temperature of 25 °C (Beanan and Strome, 1992; Mello et al., 1994). We then asked whether germline deficient animals also have suppressed the cell nonautonomous UPR^MT^ signaling **(Fig 5A)**. Indeed, we observed that germline deficient animals through the *glp-1* and *glp-4* temperature sensitive mutations suppress cell nonautonomous UPR^MT^ signaling in the neuronal PolyQ40 stress model **(Fig 5A*i-ii*)**. Interestingly, the *glp-1(e2141ts)* allele is long-lived while *glp- 4(bn2)* is not (Greer et al., 2010; TeKippe and Aballay, 2010), decoupling germline-mediated suppression of UPR^MT^ signaling from longevity mechanisms. In addition, we find that the *glp- 4(bn2)* restrictive mutation also suppresses cell nonautonomous UPR^MT^ signaling for the neuron-specific *cco-1* knockdown model **(Fig 5A*iii*)**. For a pharmacological method, we used the drug FUDR to ablate the germline through prohibiting DNA replication, especially of mitochondria in the developing germline (Tsang and Lemire, 2002) **(Fig 5B)**. We found that FUDR treatment depleted the germline of mitochondria **(Supp Fig 5A)** as well as suppressed cell nonautonomous activation the UPR^MT^ in both the neuronal PolyQ40 and *cco-1* RNAi hairpin models **(Fig 5B)**. Together, these data demonstrate that an intact germline is essential for cell nonautonomous UPR^MT^ signaling.

**Figure 5:**
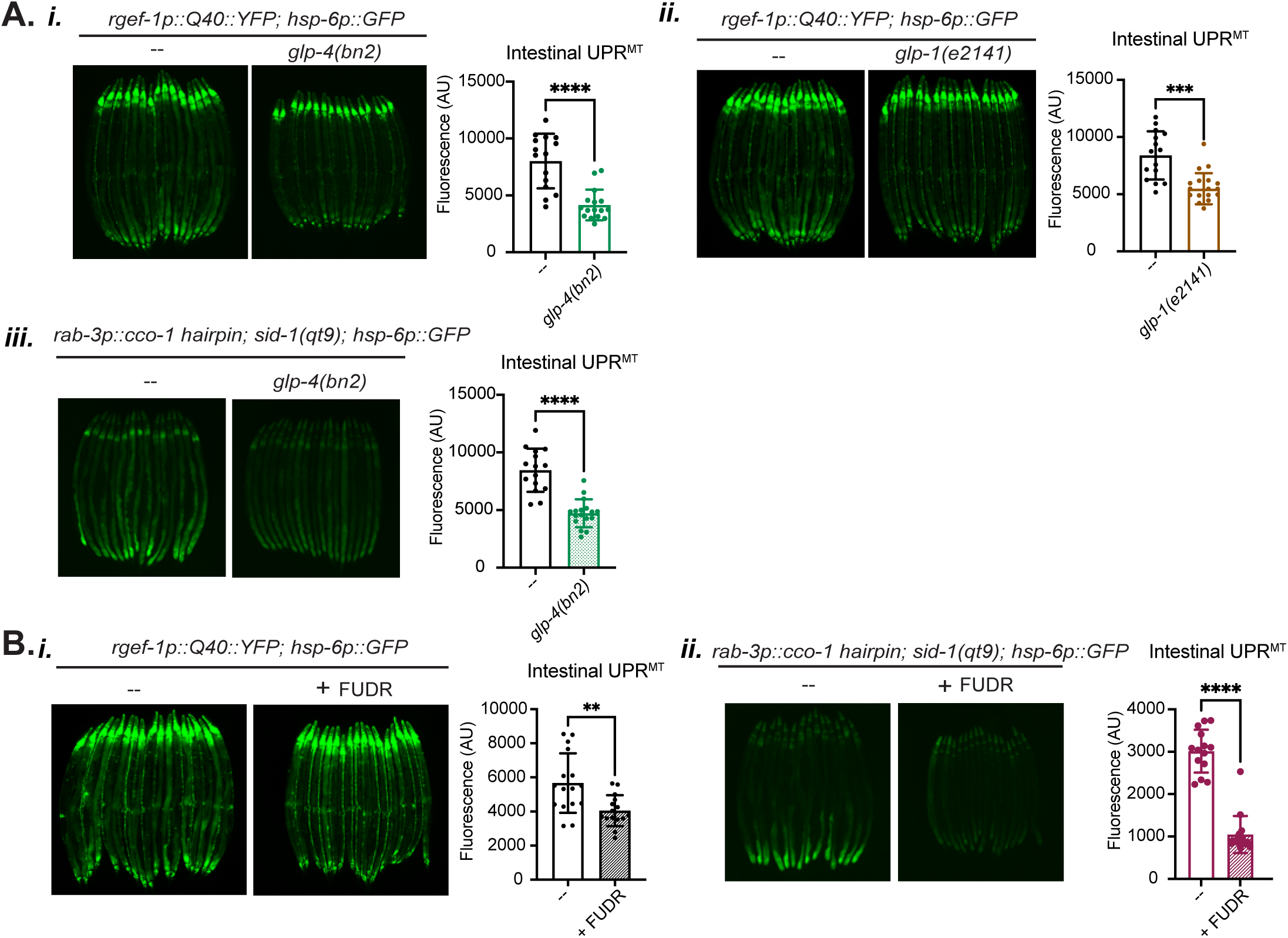
The germline, and especially germline mitochondria, is required for UPR^MT^ non- autonomous signaling. a. Comparison of fluorescence imaging and quantification of intestinal UPR^MT^ signal in *(i) glp-4(bn2)* or *(ii) glp-1(e2141)* temperature sensitive mutants, all in the *rgef-1p::Q40::YFP; hsp-6p::GFP* background, or *(iii) glp-4(bn2)* in the *sid-1(qt9); rab-3p::cco-1 hairpin; hsp- 6p::GFP* background. Experiment conducted at the restrictive temperature 25 °C to deplete the germline. Significance determined by Welch’s unpaired t-test, ****, *** p<0.0001; n = 3. b. Comparison of fluorescence imaging and quantification of intestinal UPR^MT^ signal upon FUDR treatment for germline mitochondria depletion in the *(i) rgef-1p::Q40::YFP; hsp- 6p::GFP* background or the *(ii) glp-4(bn2)* in the *sid-1(qt9); rab-3p::cco-1 hairpin; hsp- 6p::GFP* background. Significance determined by Welch’s unpaired t-test, **p = 0.0034, **** p<0.0001; n = 3.

We also tested the specificity of the germline in regulating cell nonautonomous stress response signaling, like for the UPR^ER^ or HSR. We find that FUDR treatment did not appreciably inhibit UPR^ER^ cell nonautonomous signaling in the UPR^ER^, although it did decrease the HSR model **(Supp Fig 5B)**. We observed that the FUDR-treated HSR model animals were small and sick, so the lack of HSR signal may result from additional drug interactions with FUDR treatment. We also tested whether genetic germline depletion via the *glp-4(bn2)* temperature sensitive mutation also impacted cell nonautonomous signaling in the UPR^ER^ or HSR models. While we found the signaling models to be more variable at the 25 °C restrictive temperature, we did not see appreciable reduction in cell nonautonomous signaling like for the UPR^MT^ **(Supp Fig 5C)**. Overall, these results suggest that depletion of the germline tissue or germline mitochondria specifically inhibits cell nonautonomous UPR^MT^ stress signaling between neurons and the intestine.

### *ucr-2.3* mediates cell nonautonomous UPR^MT^ downstream of established neuronal factors in UPR^MT^ signaling

Having identified the critical role of germline mitochondria and germline-acting *ucr-2.3* in mediating UPR^MT^ signaling between neurons and the intestine, we next asked whether germline *ucr-2.3* functioned downstream of previously identified neuronal mitokine signaling factors (Berendzen et al., 2016; Merkwirth et al., 2016; Zhang et al., 2018). For instance, neuronal overexpression of the WNT ligand *egl-20* and the histone demethylase *jmjd-1.2* can activate intestinal UPR^MT^ even in the absence of neuronal mitochondrial stress **(Fig 6A)** (Merkwirth et al., 2016; Zhang et al., 2018). We find that loss of *ucr-2.3* suppressed intestinal UPR^MT^ activation even in the presence of neuronal over-expression of *egl-20* and *jmjd-1.2* **(Fig 6B-C)**. This brings further evidence placing the role of *ucr-2.3* in the germline downstream of these neuronal factors to mediate cell non-autonomous UPR^MT^ signaling. Additionally, we find that germline RNAi-specific knockdown of UPR^MT^ regulators, such as *atfs-1*, *egl-20*, and *jmjd-1.2,* do not suppress cell non- autonomous UPR^MT^ signaling **(Supp Fig 6)**, suggesting that the germline has an independent role from these other neuronal factors for mediating cell nonautonomous UPR^MT^.

**Figure 6:**
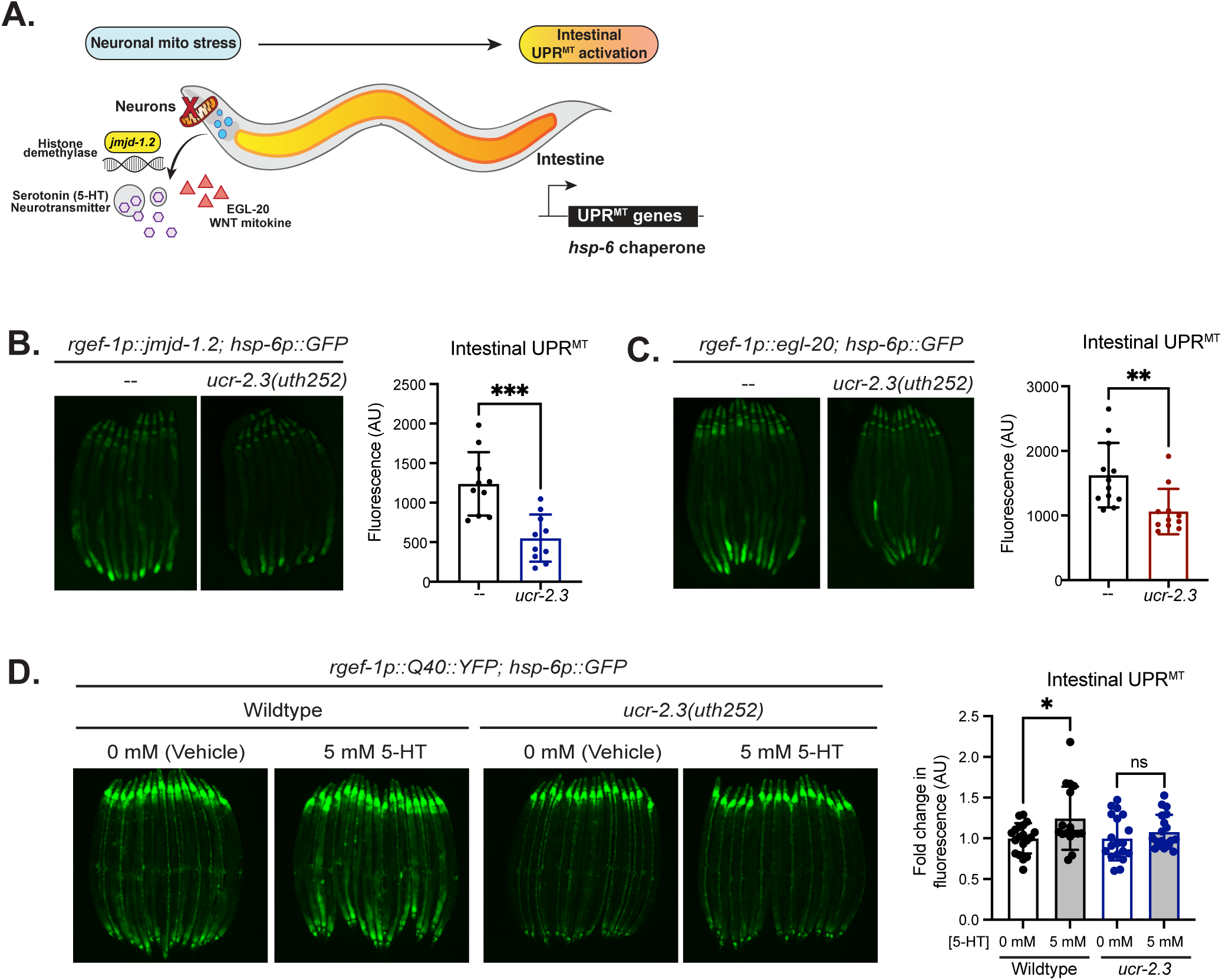
*ucr-2.3* operates downstream of established neuronal factors in mediating cell nonautonomous UPR^MT^. a. Schematic of known neuronal molecular players that mediate cell nonautonomous UPR^MT^ signaling between neurons and the intestine in *C. elegans*. b. Fluorescence imaging comparison and quantification of intestinal UPR^MT^ signal in the *rgef-1p::jmjd-1.2; hsp-6p::GFP* strain with and without the *ucr-2.3(uth252)* mutation. Significance determined by Welch’s unpaired t-test, ***p=0.0004; n = 3. c. Fluorescence imaging comparison and quantification of intestinal UPR^MT^ signal in the *rgef-1p::egl-20; hsp-6p::GFP* strain with and without the *ucr-2.3(uth252)* mutation Significance determined by Welch’s unpaired t-test, **p=0.0052; n = 3. d. Fluorescence imaging comparison of intestinal UPR^MT^ signal for exogenous addition of serotonin (5-HT) to *rgef-1p::Q40::YFP; hsp-6p::GFP* animals in the wildtype or *ucr- 2.3(uth252)* loss of function mutant. Quantification shows fold change in intestinal UPR^MT^ fluorescence upon serotonin (5-HT) addition. Significance determined by Welch’s unpaired t-test, * p = 0.0296, non-significant p value = 0.3129. n = 4

One of the factors we also considered that could signal to the germline was serotonin, a neurotransmitter that is released by dense core vesicles and mediates cell nonautonomous activation of intestinal UPR^MT^ upon neuronal mitochondrial stress (Berendzen et al., 2016). We directly tested whether *ucr-2.3* acted downstream of serotonin signaling by testing whether exogenous serotonin addition could rescue cell nonautonomous UPR^MT^ signaling between neurons and the intestine. While we found that exogenous addition of serotonin (5-HT) could increase intestinal UPR^MT^ signaling in the wildtype animal, loss of function in *ucr-2.3* prevented serotonin-mediated activation of intestinal UPR^MT^ **(Fig 6D)**. Therefore, germline *ucr-2.3* functions not only downstream of *egl-20* and *jmjd-1.2*, but also downstream of serotonin signaling pathways to mediate intestinal UPR^MT^ activation upon neuronal mitochondrial stress.

### Germline mitochondria regulate intestinal UPR^MT^ activation through altering lipid metabolic and transport pathways

Although our data indicate the *ucr-2.3* works downstream of the neuronal factors in regulating intestinal UPR^MT^ activation, our data do not rule out the possibility that *ucr-2.3* works in parallel with the neuronal factors to coordinate UPR^MT^ induction in peripheral cell types. In this scenario, a signal might be sent from the germline to work with WNT signaling in peripheral cells to specifically induce the UPR^MT^ and not developmental programs mediated by canonical WNT signaling.

To identify what signals could be sent from the germline to regulate UPR^MT^ induction in the intestine, we considered existing signaling models between the germline and the somatic tissues, such as through lipid metabolism, steroid hormones, or piRNAs (Berman and Kenyon, 2006; Goudeau et al., 2011; Han et al., 2017; Lapierre et al., 2011; Shi and Murphy, 2023; Wang et al., 2008). We were particularly motivated to explore a lipid-based germline signaling model since one of our initial observations of the *ucr-2.3* mutant was that it had drastically higher levels of intestinal fat **(Fig 7A, Supp Fig 7A)**. Since *ucr-2.3* mutant animals have compromised germline mitochondrial and tissue integrity, this is in alignment with reports by others that germline deficient animals also have higher intestinal fat levels **(Supp Fig 7B)** (O’Rourke et al., 2009). In addition, from an RNA-seq analysis comparing *ucr-2.3(uth252); rgef-1p::Q40::YFP* mutant animals to *rgef-1p::Q40::YFP* alone, we found that many up-regulated transcripts in the *ucr- 2.3(uth252)* mutant animals were highly enriched in GO terms for metabolic processes, especially those involving fatty acids and lipids **(Fig 7B)**. Therefore, we tested whether compromised germline mitochondrial integrity resulted in suppressed intestinal UPR^MT^ activation through altering lipid metabolic pathways in the intestine. We conducted a targeted RNAi screen of fatty acid and lipid metabolic genes corresponding to the up-regulated GO terms in the *ucr-2.3(uth252)* mutant. Surprisingly, we found that knockdown of *fat-2*, a Δ12 desaturase enzyme that converts oleic acid to linoelic acid in the first step of polyunsaturated fatty acid (PUFA) synthesis in *C. elegans* (Watts and Browse, 2002; Zhou et al., 2011), rescued nonautonomous UPR^MT^ signaling in the *ucr-2.3(uth252)* mutant **(Fig 7C)**.

**Figure 7:**
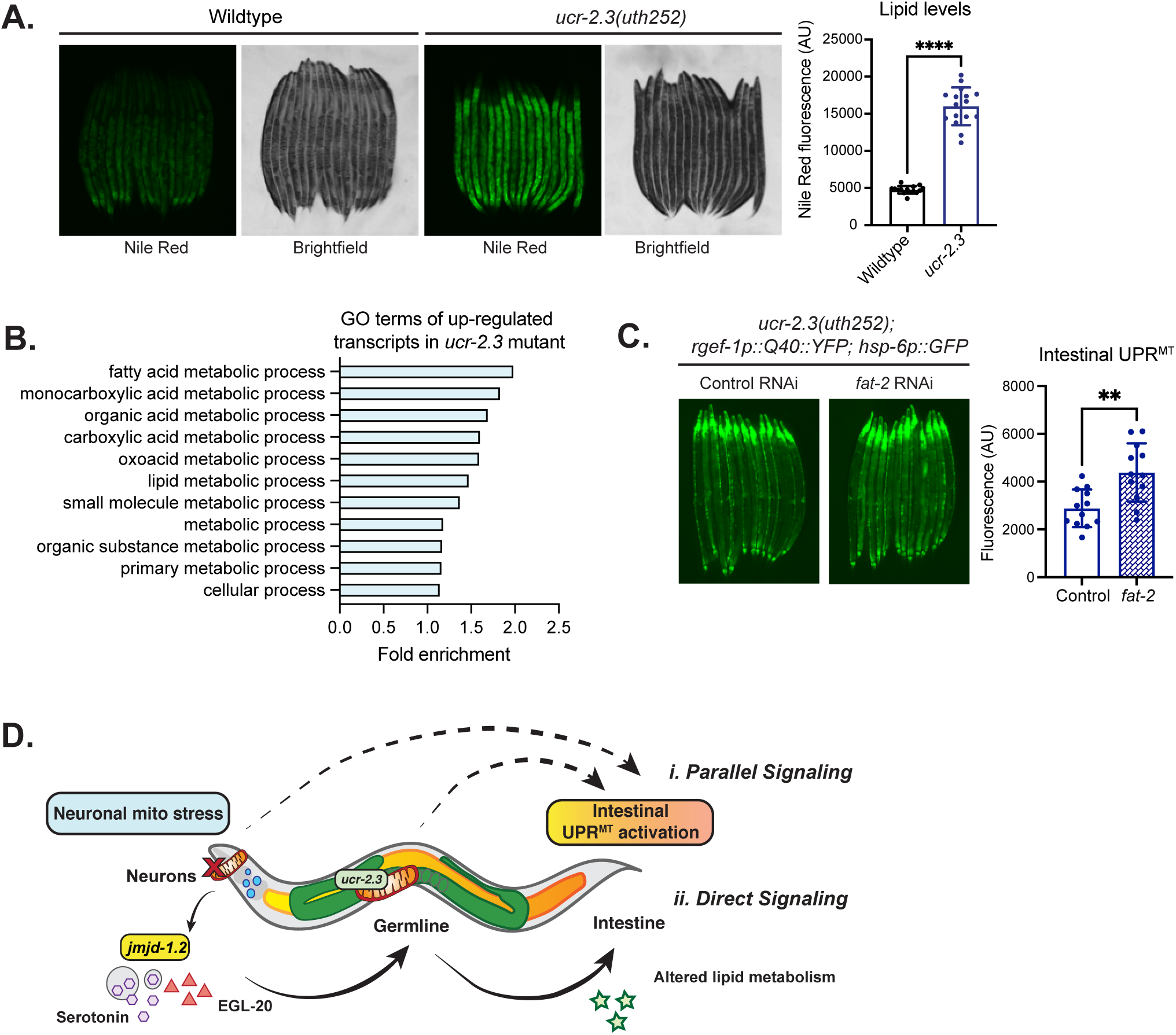
The germline regulates intestinal UPR^MT^ activation in the intestine through alteration of lipid metabolic pathways. a. Comparison of lipid levels and quantification of Nile Red fluorescence between wildtype and the *ucr-2.3(uth252)* mutant. Significance determined by Welch’s unpaired t-test, ****p <0.0001; n = 3. b. Enriched GO terms (biological process) in up-regulated transcripts in the *ucr-2.3(uth252); rgef-1p::Q40::YFP* mutant compared to *rgef-1p::Q40::YFP* alone. Shown are Fold enrichment values determined by the PANTHER statistical overrepresentation test. c. Fluorescence comparison of *ucr-2.3(uth252); rgef-1p::Q40::YFP; hsp-6p::GFP* animals on control or *fat-2* RNAi. Significance determined by Welch’s unpaired t-test, **p = 0.0020; n = 2. d. Proposed model for how germline mitochondria mediate neuron-to-intestine UPR^MT^ stress signaling. *(i)* Parallel Signaling pathway (Dashed lines): Neurons can communicate to the intestine, or the germline. Simultaneously, the germline acts as a master switch to control “ON”/”OFF” ability for the UPR^MT^ to be activated in the intestine, regardless of presence of other signals. *(ii)* Direct Signaling pathway (Solid lines): Neurons directly communicate to the germline, which processes signals possibly through mitochondrial activity, then sends a signal to the intestine to regulate UPR^MT^ activation.

We also considered pathways involving direct transfer of lipid molecules between the germline and the intestine, such as the *vit* family of vitellogenins are lipoproteins which transport phospholipids, cholesterol, and other nutrients from the intestine to the germline (Perez and Lehner, 2019) and are the evolutionary ancestor of human apolipoprotein B (APOB) (Babin et al., 1999). We find that the *vit* family genes (*vit-1, 2, 3, 5, 6*) are also transcriptionally up-regulated in the *ucr-2.3* mutant RNA-seq dataset (mean log2(FC) = 0.669, SD = 0.191). Additionally, we find that RNAi knockdown of the *vit* family genes, which redistributes lipids and increases lysosomal lipolysis in the intestine (Seah et al., 2016), rescued intestinal UPR^MT^ activation in the *ucr-2.3* mutant **(Supp Fig 7C)**.

Overall, we find that *ucr-2.3* mutant animals have up-regulated intestinal lipid levels and metabolic genes, and manipulating lipid synthesis and transport pathways in the intestine can rescue non-autonomous UPR^MT^ activation in the *ucr-2.3* mutant. This suggests a mode of lipid-based signaling between germline mitochondria and the intestine, whereby germline mitochondrial quality may be communicated to the intestine through lipid synthesis and transport pathways which subsequently regulates UPR^MT^ activation. Indeed, previous work had shown that mitochondrial stress signaling can be also regulated by ceramide and cardiolipin lipids levels (Kim et al., 2016), suggesting a general principle of mitochondrial stress regulation by lipids.

Collectively our data propose a model in which the germline, through germline mitochondria, integrates mitokine stress signals from the nervous system to regulate UPR^MT^ in peripheral tissues, such as the intestine **(Fig 7D)**. This highlights the role of the germline as a signaling tissue that actively communicates with the soma regarding mitochondrial stress to influence organismal stress and metabolic phenotypes. We believe the evolutionary basis for this neuron and germline signaling circuit may be to conserve overall organismal resources in the somatic tissues. The need to activate energy-intensive, protective stress responses should only be executed in the presence of a functioning germline to preserve progeny production and continuation of the immortal germline. Without a functional germline, there would be no need to activate these energy-intensive stress responses, and the UPR^MT^ is shut off to direct resources elsewhere. On a broader level, we believe this signaling circuit has implications for how germ cells and germline mitochondrial quality may be influenced by and respond to stresses occurring in the somatic environment, such as with aging, metabolic disease, or infection.

## Discussion

In this study, we have identified the importance of the germline, and particularly of germline mitochondria, in mediating cell nonautonomous communication of mitochondrial stress from the nervous system to peripheral cells. Using a genome-wide mutagenesis screen, we identified the mitochondrial gene *ucr-2.3* as a novel regulator of cell nonautonomous UPR^MT^ signaling between neurons and the intestine in *C. elegans*. Surprisingly, we find that *ucr-2.3* mediates cell nonautonomous UPR^MT^ signaling through its activity in the germline tissue. Loss of *ucr-2.3* activity in the germline compromises germline mitochondrial health, germline integrity, and suppresses neuronal-to-intestinal communication of mitochondrial stress signals. Furthermore, we find that *ucr-2.3* works downstream and independently of other identified neuronal mediators of cell nonautonomous UPR^MT^ signaling such as serotonin and *egl-20*/WNT mechanisms. We also observe that *ucr-2.3* and germline tissue integrity is specific to UPR^MT^ cell nonautonomous signaling and is not a primary mediator for other forms of cell nonautonomous communication, such as the UPR^ER^ or cytosolic HSR. This suggests that there is a specific relationship between germline mitochondrial integrity and inter-tissue mitochondrial stress signaling in the somatic tissues. Finally, we find that *ucr-2.3* activity in the germline may communicate to the intestine through alteration of lipid metabolism pathways to regulate UPR^MT^ activation. We propose that this germline-to-intestine signal could also direct WNT signaling factors from their role in developmental programs to regulating UPR^MT^ activation (Zhang et al., 2018).

Overall, our data highlight the role of the germline tissue in mediating cell nonautonomous mitochondrial stress signaling and specifically a specialized role for germline mitochondria. Our results provide additional context for understanding the communication between germline and somatic tissues in regulating organismal health, especially regarding mitochondrial quality. While often regarded as two separate entities, the soma and the germline must engage in constant communication with each other to ensure overall organismal health (Conine and Rando, 2022). This can come in the form of small RNAs, nutrients, and other growth or stress cues that allow the germline to adapt to sources of environmental or somatic stress and, conversely, allow the somatic tissues to respond to changes in germline integrity. Several examples of soma-to-germline communication pathways have been described in *C. elegans* (Calculli et al., 2021; Curran et al., 2009; Shi and Murphy, 2023). Our work here illustrates a germline mitochondrial-specific communication pathway that involves reshuffling of lipid transport and metabolism pathways in the intestine in response to mitokine stress signals from neurons.

Our findings suggest two models for how the germline could mediate neuronal-to-intestinal UPR^MT^ signaling **(Fig 7D)**. In one model (“Parallel Signaling”), the germline acts as an ON/OFF switch for intestinal UPR^MT^ activation. In this model, peripheral activation of the UPR^MT^ requires signals from the nervous system (e.g., WNT and serotonin) *and* a signal from an intact, healthy germline. In another model (“Direct Signaling”), the germline could act as an intermediary tissue, wherein the germline receives signals secreted by neurons experiencing mitochondrial stress, processes these signals – perhaps through mitochondrial *ucr-2.3* activity – then emits an additional signal to the intestine to regulate UPR^MT^ activation. In both models, the germline may regulate intestinal activation of UPR^MT^ signaling through lipid transport and synthesis pathways.

Based on our findings, we cannot disprove either the Parallel or Direct Signaling model, nor have we shown direct evidence that germline mitochondria can receive signals from neurons. However, regarding the possibility of a Direct Signaling model, we do observe that germline mitochondrial signaling operates downstream of serotonin **(Fig 6D)**. In addition, it had been previously shown that in the same neuronal polyQ40 model of mitochondrial stress in *C. elegans*, neuronal mitochondrial stress elevated mtDNA levels in germline mitochondria, which could be transgenerationally inherited (Q. Zhang et al., 2021). Furthermore, components of the WNT signaling pathway, such as the beta-catenin *bar-1,* were also shown to directly regulate oocyte mitochondria mass and mtDNA levels (Q. Zhang et al., 2021). Serotonin signaling has also been shown to directly improve oocyte quality in *C. elegans* and *D. melanogaster* (Aprison et al., 2023), suggesting that a Direct Signaling route between neurons and germ tissue through serotonin-based signaling could exist.

The mechanism for how *ucr-2.3* activity in the germline controls cell nonautonomous UPR^MT^ remains to be explored. One possibility is that loss of *ucr-2.3* function leads to compromised mitochondrial function in the germ cells **(Fig 2C**, **Fig 4D, Supp Fig 5A)**, leading to a general depletion of germ cell proliferation and resulting germline tissue loss. This is reflected in the very small brood size of *ucr-2.3* mutant animals **(Fig 4C)**. In this scenario, the loss of functional *ucr-2.3* compromises general germline integrity, leading to a loss of cell nonautonomous signaling. Indeed, we believe the germline-specific expression of *ucr-2.3* is why only mutations in *ucr-2.3* mutants were uncovered in our screen, as opposed to its more ubiquitously expressed (and germline excluded) paralogs *ucr-2.1/2.2*.

Beyond its role in the electron transport chain, there could also be a specific activity related to *ucr-2.3* in generating a signal either required for germline mitochondrial integrity or germline-to-intestine UPR^MT^ signaling. One possibility we considered is the predicted role of *ucr-2.3* as a protease, as its protein sequence is predicted to belong to the M16 metallopeptidase family, which includes mitochondrial processing peptidases. Indeed, the mammalian ortholog of *ucr-2.3*, UQCRC2, is also predicted to have MPP-like (mitochondrial peptide processing) protease activity in combination with UQCRC1 to cleave off the N-terminal MTS of UQCRFS1 (the Rieske Fe-S protein) in the final steps of complex III assembly (Deng et al., 2001, 1998; Fernandez-Vizarra and Zeviani, 2018). Whether *C. elegans* UCR-2.3 could also have an MPP-like role in complex III assembly in germline mitochondria or have an additional peptidase role to generate peptide-like signals in the context of mitochondrial stress are interesting hypotheses that should be further explored. While it is intriguing to speculate that *ucr-2.3* is specific for the germline signal, additional germline restricted ETC genes, such as *nduf-2.2* were also required **(Fig 4G)**. Therefore, it is also possible that *nduf-2.2* and *ucr-2.3* work together to generate a germline specific signal, or more likely, general mitochondrial integrity within the germline is the predominant source of the signal to the intestine. While each possibility cannot be excluded, it is intriguing to speculate that evolution has opted to restrict expression of a few, key ETC genes within the germline to help coordinate signals to the soma. Whether mitochondrial ETC isoforms with tissue-specific expression also exist in humans should also be explored.

While several reports of germline deficient animals have increased stress resistance for the cytosolic HSR or lifespan phenotypes (Hsin and Kenyon, 1999; Labbadia and Morimoto, 2015), we find the opposite for mitochondrial stress responses; specifically, that germline deficiency shuts down UPR^MT^ signaling. We believe this specificity in germ cell regulation of mitochondrial stress signaling may result from a special requirement of mitochondrial quality in the germline. Because of the high abundance of mitochondria in germ cells and their important role in germ cell maturation (Van Blerkom, 2011; N. Zhang et al., 2021), the integrity of the germline and its consequential communication with other tissues may have a heavy reliance on mitochondrial quality. Indeed, oocytes in *C. elegans*, *Xenopus*, and humans employ specialized mechanisms to maintain mitochondrial quality and ensure integrity of the reproductive tissues (Colnaghi et al., 2021; Cota and Murphy, 2021; Rodríguez-Nuevo et al., 2022). For instance, germline mitochondria demonstrate a higher protein import capacity compared to somatic mitochondria in *C. elegans* (Xin et al., 2022). Our results also highlight the importance of mitochondrial quality in germ tissue and suggest the possibility that integrity of germline may also regulate stress signaling pathways between the germline and somatic tissues. It will be important to understand whether these germline-somatic mitochondrial stress signaling pathways also are relevant for other forms of somatic stress besides proteotoxic PolyQ40 aggregation, such as obesity or infection, and what implications stresses occurring in the somatic tissues may have on germline mitochondrial quality, fertility or reproductive tissue health, and lipid metabolism.

Finally, it is intriguing to consider our results in the context of the “Disposable Soma Theory” which contends that there is an energetic trade-off between somatic maintenance and reproductive health. By this theory, animals lacking a germline should be able to invest more energetic resources to health maintenance in the somatic tissues, leading to increased longevity and stress resistance. Our observations provide another layer of complexity to this theory by contending that there may be an energetic triage occurring in response to loss of germline tissue. While loss of germline tissue or *ucr-2.3* may lead to the up-regulation of certain stress responses, it leads to an intestinal UPR^MT^ shutdown. This may operate through signaling mechanisms specific to mitochondrial health or mitochondrial substrate requirements for metabolism no longer necessary in an infertile organism. Thus, this intestinal UPR^MT^ shut off could be an adaptive mechanism to conserve organismal resources but have additional consequences on fat metabolism in somatic tissues. Whether infertility or compromised reproductive tissues health may have similar consequences in mitochondrial or metabolic states in humans is of great future interest, especially given existing connections between obesity and sterility/infertility in humans (Carrageta et al., 2019; Ozcan Dag and Dilbaz, 2015; Whon et al., 2021).

## Acknowledgements

We thank all members of the Dillin lab, Dr. Jeremy Chang, Dr. Veerle Rottiers, and Dr. Malene Hansen for their feedback on the project. We acknowledge support by the Howard Hughes Medical Institute (J.K.D., C.G.M., L.J., A.D.) and the National Institutes of Health (A.D., B.M.W.). K.S. is supported by a K99/R00 Pathway to Independence Award (K99AG071935). C.K.T. is supported by a NIH F32 Postdoctoral Fellowship (5F32AG069388- 03). H.Z. is supported by the Larry L. Hillblom Foundation Postdoctoral Fellowship (2020-A-018- FEL) and the Glenn Foundation for Medical Research Postdoctoral Fellowship.

---

**Supplementary Figure 1:**
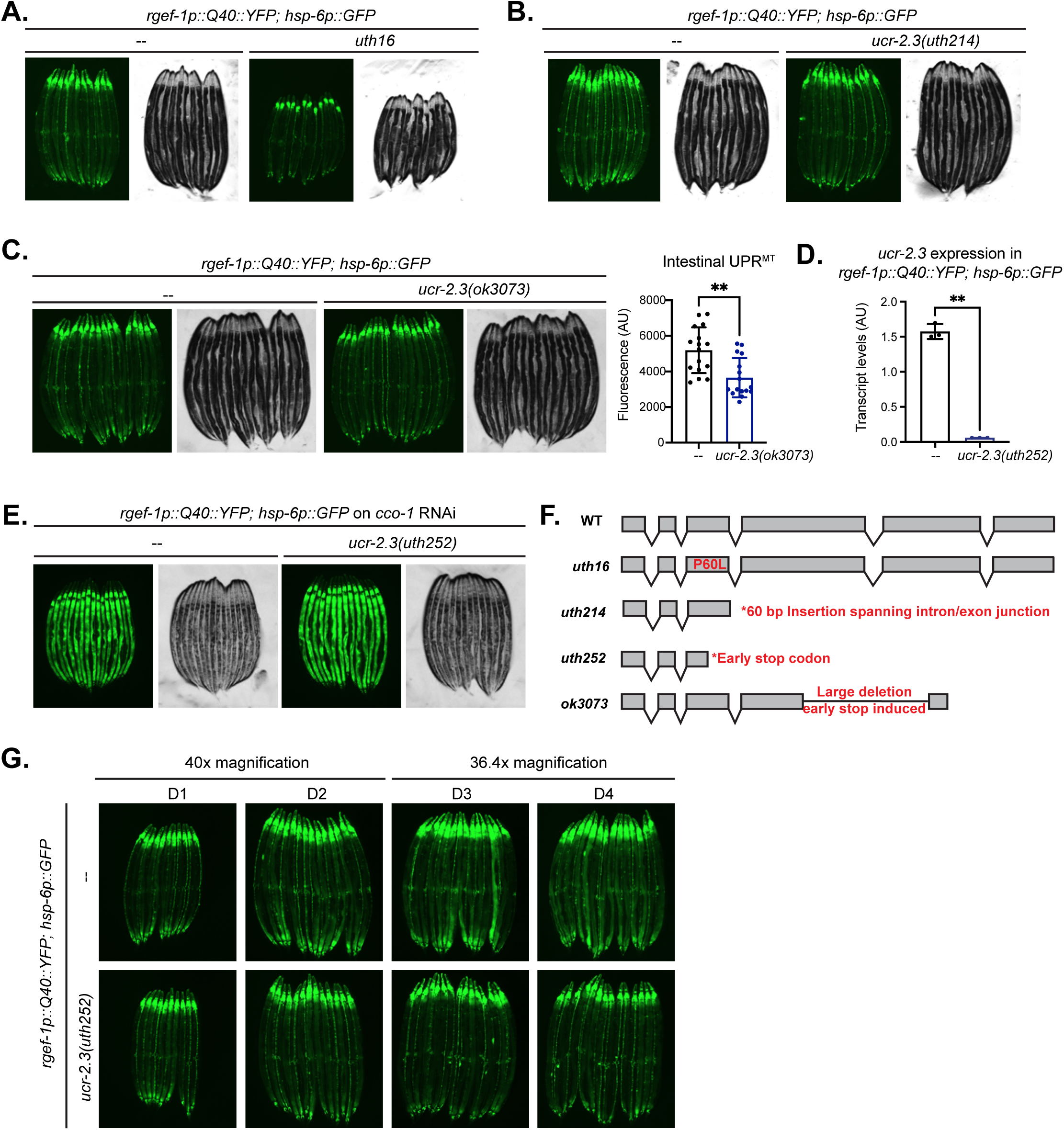
Additional genetic evidence supporting loss of function mutations in *ucr-2.3* suppress cell nonautonomous UPR^MT^ signaling in *C. elegans*. a. Fluorescence imaging comparison of the wildtype UPR^MT^ cell nonautonomous reporter strain *rgef-1p::Q40::YFP;hsp-6p::GFP* to the EMS mutant obtained in the suppressor screen *uth16*. n> 3. b. Fluorescence imaging comparison of a loss of function mutation in *ucr-2.3(uth214)* generated by CRISPR/Cas9 genome editing to a non-edited, wildtype strain of *rgef- 1p::Q40::YFP;hsp-6p::GFP*. n > 3. c. Fluorescence imaging comparison and quantification of a large deletion mutation in *ucr-2.3* compared to a wildtype strain of *rgef-1p::Q40::YFP;hsp-6p::GFP*. Significance determined by Welch’s unpaired t-test, p = 0.0010; n > 3. d. qPCR comparison of *ucr-2.3* expression levels in wildtype and *ucr-2.3(uth252)* mutant animals in the *rgef-1p::Q40::YFP; hsp-6p::GFP* genetic background. Transcript levels normalized by *rpl-32*. Significance determined by Welch’s unpaired t-test, **p=0.0016. e. Fluorescence imaging showing autonomous UPR^MT^ activation comparing wildtype and *ucr-2.3(uth252)* animals fed *cco-1* RNAi in the *rgef-1p::Q40::YFP; hsp-6p::GFP* genetic background. n > 3. f. Schematic describing genetic changes in all shown mutant alleles of *ucr-2.3*. g. Fluorescence comparison of intestinal UPR^MT^ signal in wildtype and *ucr-2.3(uth252)* mutant animals over adult aging (Day 1 – 4 of adulthood). n > 3.

**Supplementary Figure 2:**
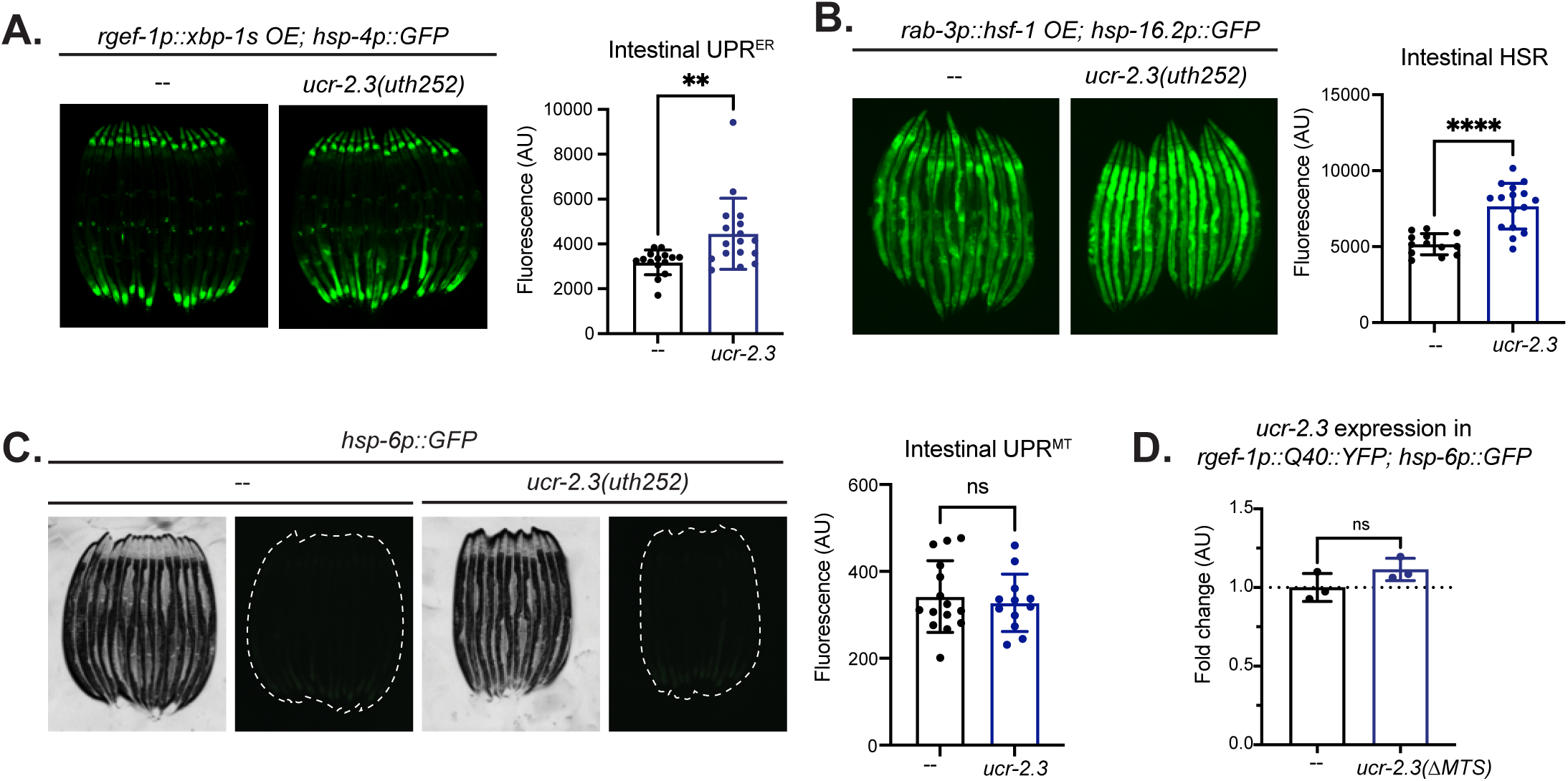
*ucr-2.3* loss of function does not suppress UPR^ER^ or cytosolic HSR cell nonautonomous signaling. a. Fluorescence imaging comparison of the cell nonautonomous UPR^ER^ reporter strain (*rgef- 1p::xbp-1s OE; hsp-4p::GFP*) with and without the *ucr-2.3(uth252)* loss of function mutation. Significance determined by Welch’s unpaired t-test, ** p = 0.0055; n = 3. b. Fluorescence imaging comparison of the cell nonautonomous HSR reporter strain (*rab- 3p::hsf-1 OE; hsp-16p::GFP)* with and without the *ucr-2.3(uth252)* loss of function mutation. Significance determined by Welch’s unpaired t-test, **** p < 0.0001; n = 2. c. Fluorescence imaging showing UPR^MT^ activation in the absence of mitochondrial stress (basal condition). Shown are wildtype and *ucr-2.3(uth252)* animals fed control RNAi in the *hsp-6p::GFP* UPR^MT^ reporter genetic background. Significance determined by Welch’s unpaired t-test, p = 0.6179; n = 3. d. qPCR comparison of *ucr-2.3* expression levels in wildtype and *ucr-2.3(11MTS)* mutant animals in the *rgef-1p::Q40::YFP; hsp-6p::GFP* genetic background. Transcript levels normalized by *rpl-32*. Significance determined by Welch’s unpaired t-test, p = 0.1603.

**Supplementary Figure 3:**
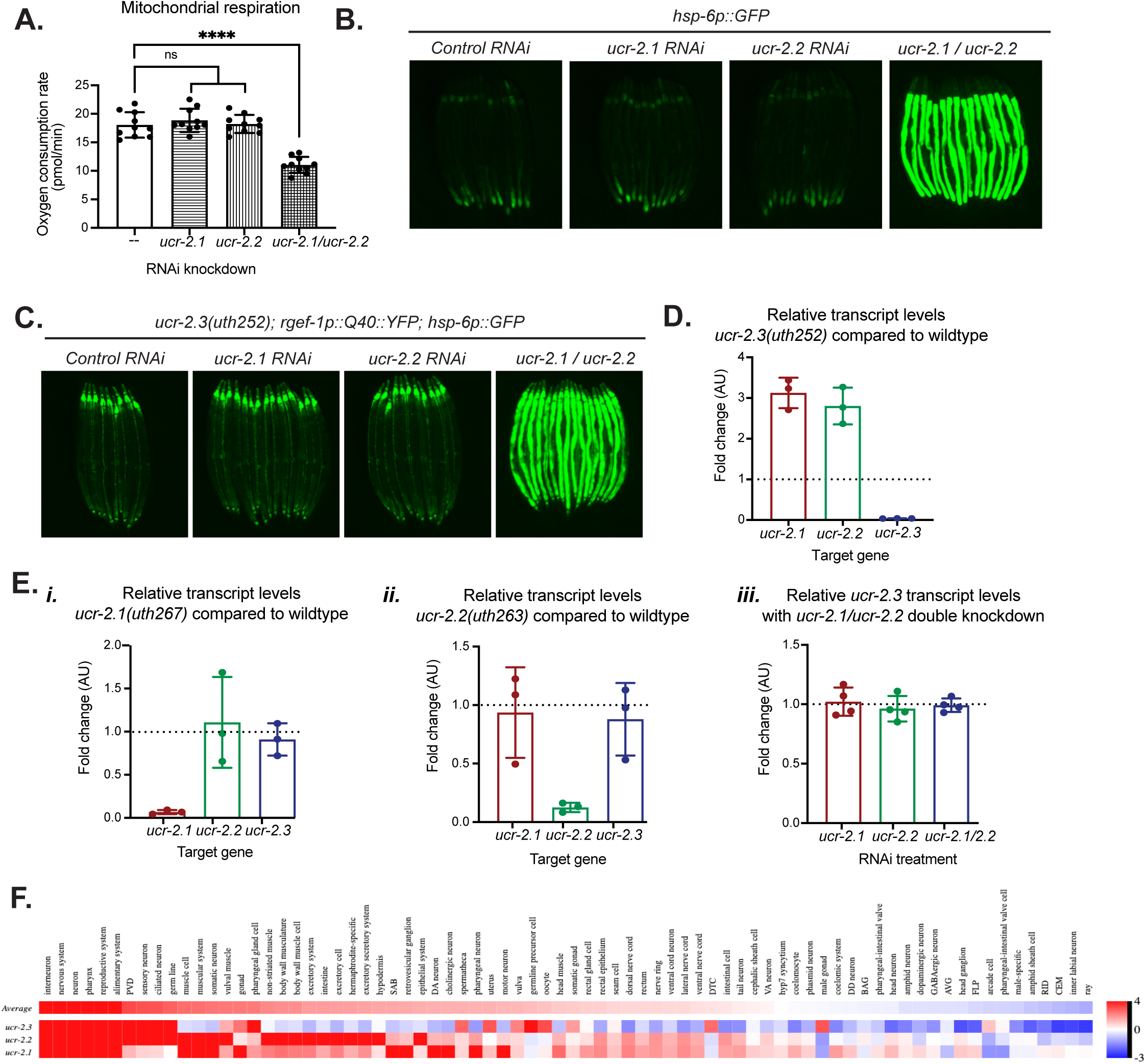
*ucr-2.1* and *ucr-2.2* are functionally redundant and display similar tissue expression patterns but differ from *ucr-2.3*. a. Measurement of mitochondrial respiration (oxygen consumption rate) upon *ucr-2.1* and *ucr-2.2* single and double RNAi knockdown in the *hsp-6p::GFP* genetic background. Significance determined by Welch’s unpaired t-test. ****p<0.0001, non-significant p values > 0.4; n = 3. b. Fluorescence imaging comparison of UPR^MT^ activation from *ucr-2.1* and *ucr-2.2* single and double RNAi knockdowns in the *hsp-6p::GFP* UPR^MT^ reporter strain. n = 3. c. Fluorescence imaging comparison of UPR^MT^ activation from *ucr-2.1* and *ucr-2.2* single and double RNAi knockdowns in the *ucr-2.3(uth252); rgef-1p::Q40::YFP; hsp-6p::GFP* genetic background. n = 3. d. qPCR measurement of relative transcript levels of the *ucr-2* family genes between the *ucr- 2.3(uth252)* loss of function mutant and wildtype in the *rgef-1p::Q40::YFP; hsp-6p::GFP* genetic background. e. qPCR measurement of relative transcript levels of the *ucr-2* family genes between the *ucr- 2.1(uth267)* mutant *(i)* and *ucr-2.2(uth263)* mutant *(ii)* and wildtype with RNAi single and double knockdown of *ucr-2.1* and *ucr-2.2 (iii)* in the *rgef-1p::Q40::YFP; hsp-6p::GFP* genetic background. f. Comparison of tissue-expression profiles for the UCR-2 family genes. Plots were generated using the Worm tissue expression prediction web interface (Kaletsky et al., 2018) (https://worm.princeton.edu/).

**Supplementary Figure 4:**
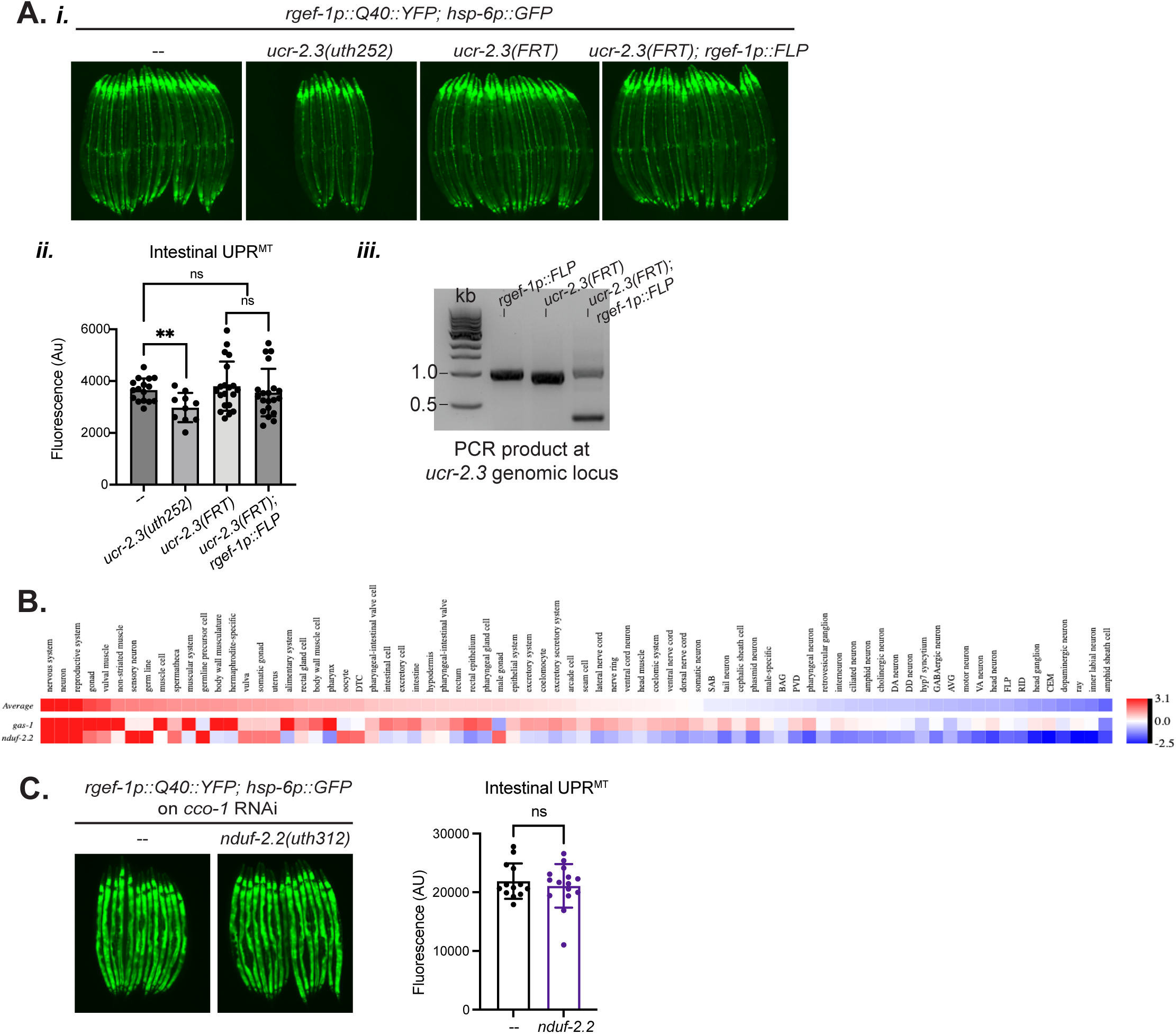
Additional genetic evidence supporting a germline-specific role for *ucr-2.3* and germline mitochondria in mediating cell nonautonomous UPR^MT^. a. *(i)* Fluorescence imaging comparison of intestinal UPR^MT^ signal between *ucr-2.3* excised only in neurons using FLP/FRT recombination (*ucr-2.3(FRT); rgef-1p::FLP*) compared to *ucr-2.3* with the FRT sites alone *ucr-2.3(FRT)*, all in the *rgef-1p::Q40::YFP; hsp-6p::GFP* genetic background. Wildtype and *ucr-2.3(uth252)* loss of function mutant displayed for comparison. *(ii)* Quantification. Significance determined by Welch’s unpaired t-test, **p=0.0052, non-significant p values > 0.4. n = 3. *(iii)* DNA gel displaying excised *ucr-2.3* upon FLP/FRT recombination by PCR genotyping the *ucr-2.3* genetic locus in each strain listed. b. Comparison of tissue-expression profiles for the *nduf-2* family genes (*gas-1/nduf-2.1*, *nduf- 2.2*). Plots were generated using the Worm tissue expression prediction web interface (Kaletsky et al., 2018) (https://worm.princeton.edu/). c. Fluorescence imaging showing autonomous UPR^MT^ activation comparing wildtype and *nduf-2.2(uth312)* animals fed *cco-1* RNAi in the *rgef-1p::Q40::YFP;hsp-6p::GFP* genetic background. n = 2.

**Supplementary Figure 5:**
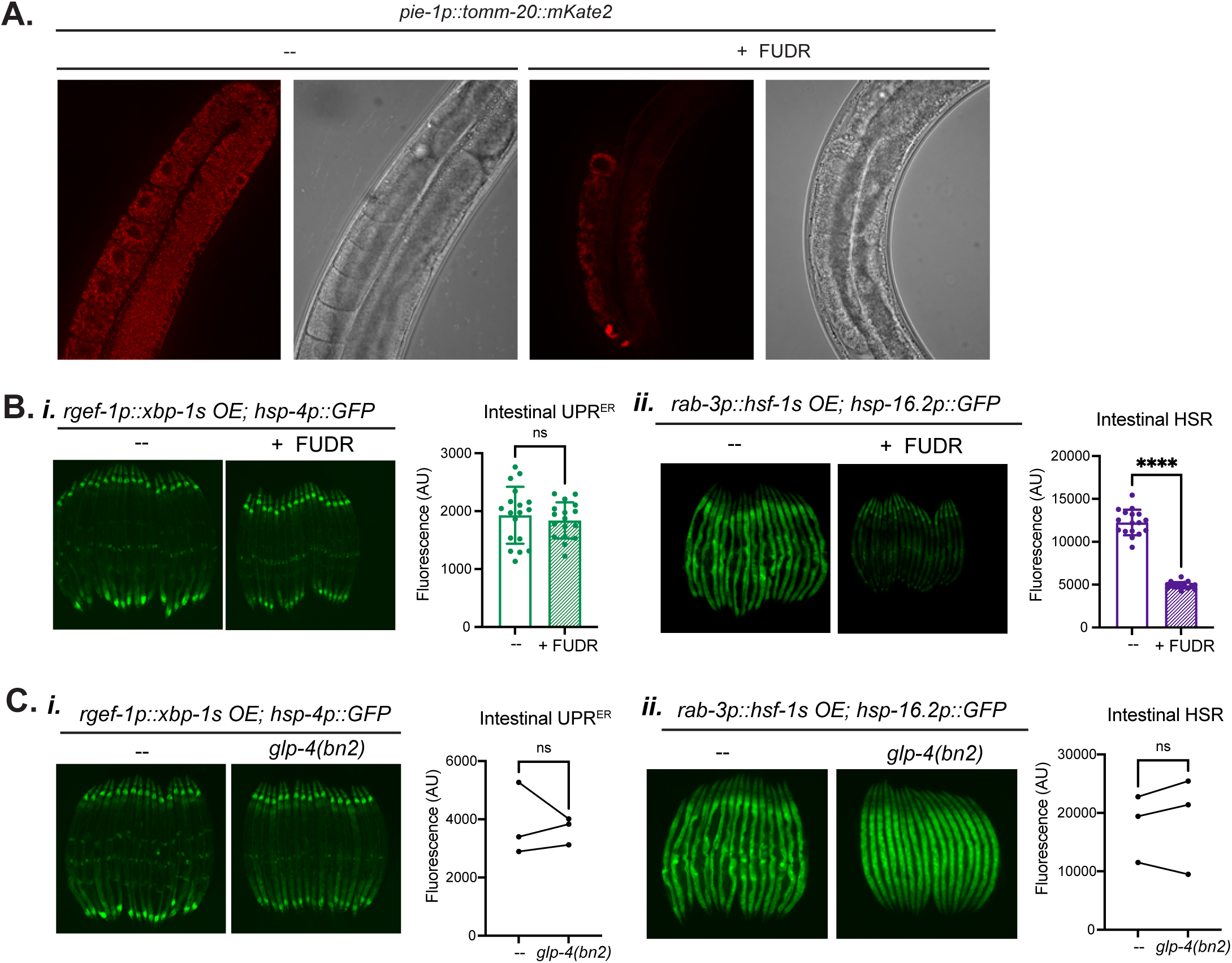
Additional evidence supporting a specific role for the germline in mediating UPR^MT^ signaling. a. Fluorescence widefield imaging of *pie-1p::tomm-20::mKate2* germline mitochondrial reporter strain with and without FUDR treatment. b. Fluorescence imaging comparison and quantification of the cell nonautonomous UPR^ER^ *(i)* and cytosolic HSR *(ii)* reporter strains upon germline depletion by FUDR treatment. Significance determined by Welch’s unpaired t-test, *(i)* p = 0.5250. *(ii)* **** p < 0.0001. n = 3. c. Fluorescence imaging comparison and quantification of the cell nonautonomous UPR^ER^ and HSR reporter strains upon genetic germline depletion by the *glp-4(bn2)* temperature sensitive mutation at the restrictive temperature 25 °C. Images shown are representative of three independent experiments. Quantification displays average fluorescence values for each strain from three independent experiments. Significance determined by Welch’s paired t-test across the three independent experiments, *(i)* p = 0.7477; *(ii)* p = 0.6100.

**Supplementary Figure 6:**
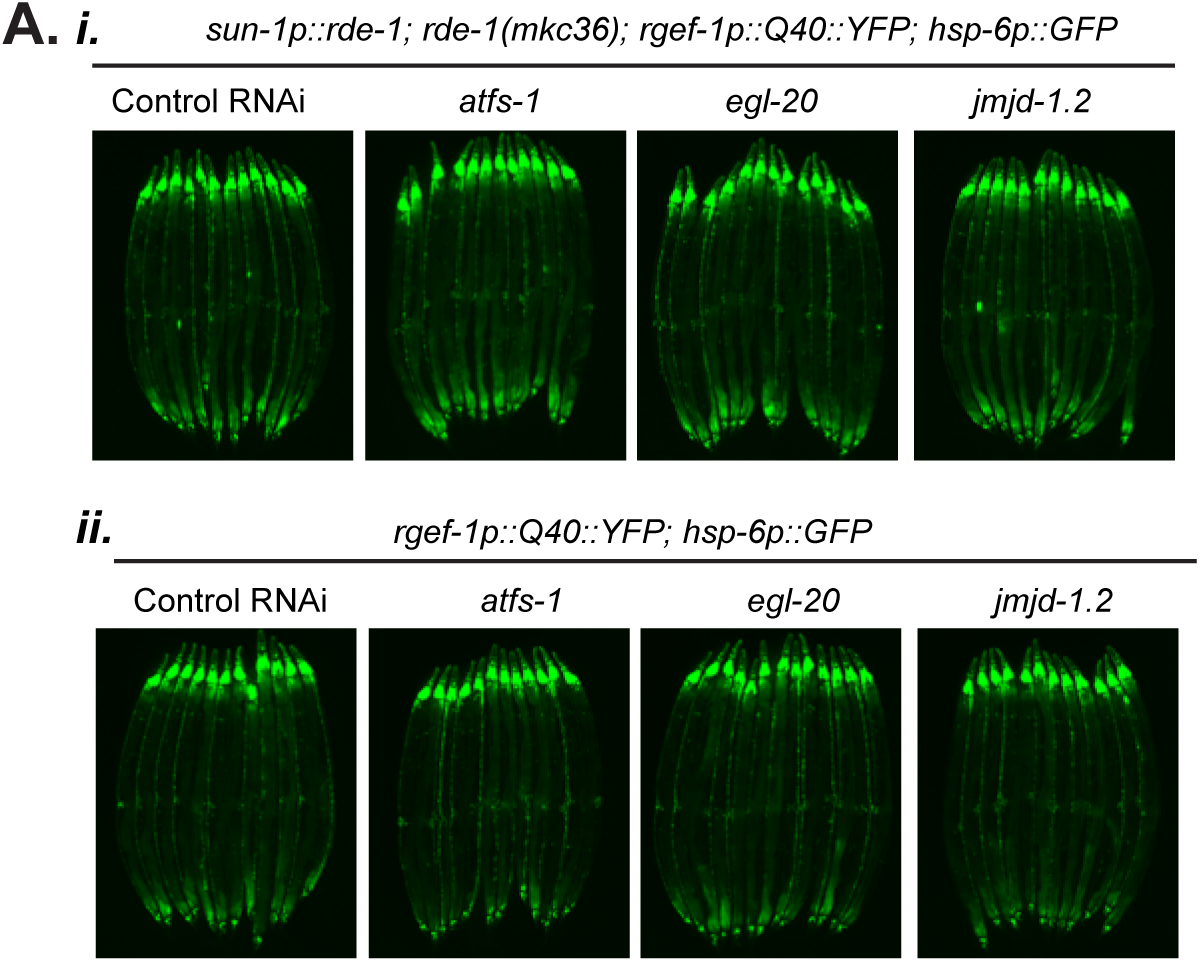
Germline-specific knockdown of known UPR^MT^ and mitokine factors does not suppress nonautonomous UPR^MT^ signaling. a. *(i)* Fluorescence imaging comparison of intestinal UPR^MT^ signal for RNAi knockdown of UPR^MT^ factors *atfs-1*, *egl-20*, and *jmjd-1.2* in a germline-specific RNAi strain (Zou et al., 2019) crossed to the cell nonautonomous UPR^MT^ reporter: *sun-1p::rde-1; rde-1(mkc36); rgef-1p::Q40::YFP; hsp-6p::GFP*. *(ii)* RNAi knockdown of the same UPR^MT^ factors in the standard cell nonautonomous UPR^MT^ reporter *rgef-1p::Q40::YFP; hsp-6p::GFP*. n = 3.

**Supplementary Figure 7:**
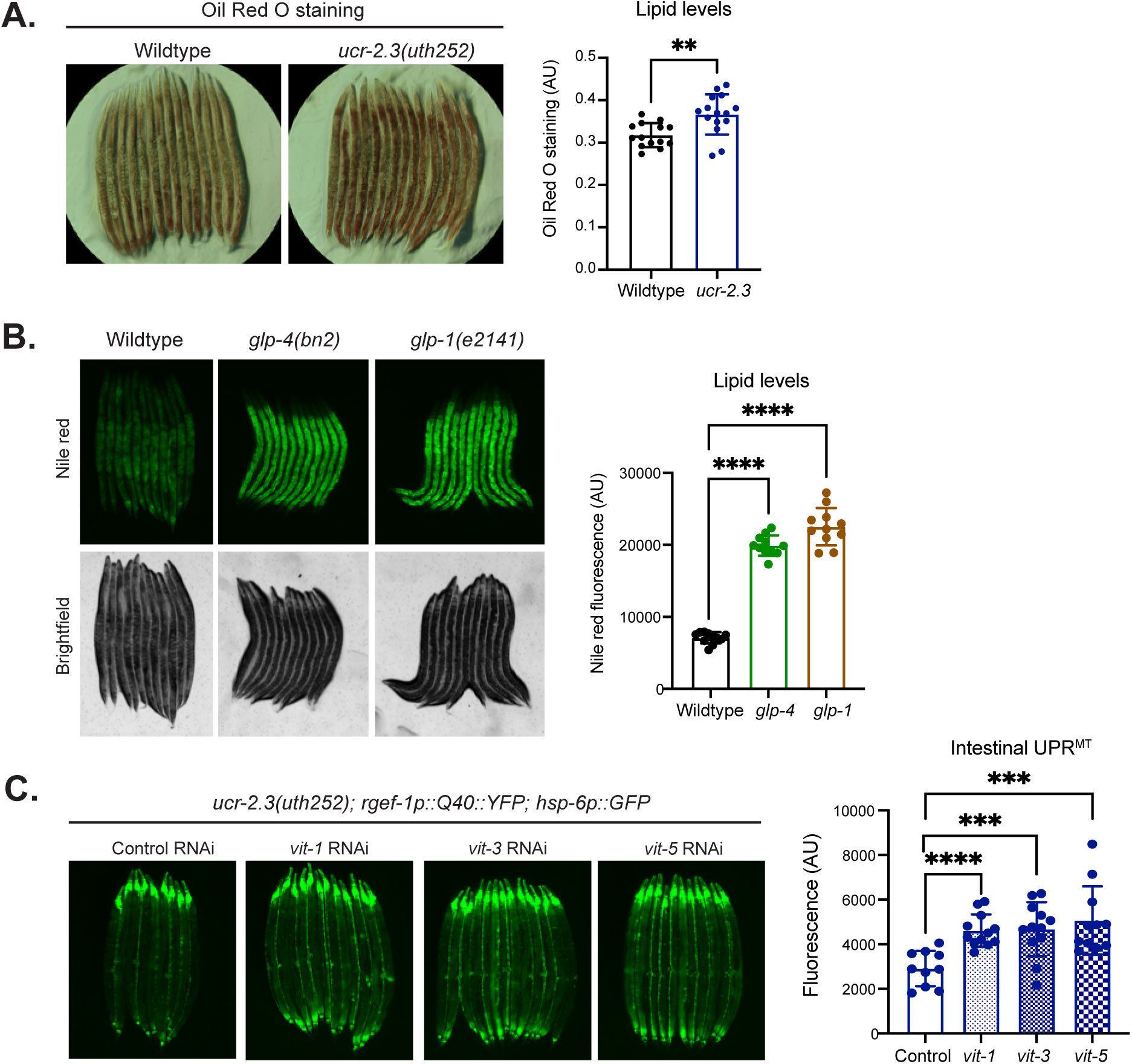
Increased intestinal fat levels in *ucr-2.3* loss of function and germline deficient animals. a. Oil Red O staining and quantification of lipid levels in wildtype or *ucr-2.3(uth252)* mutant animals. Significance determined by Welch’s unpaired t-test, ** p = 0.0185; n = 2. b. Comparison of lipid levels and quantification of Nile Red fluorescence between wildtype and germline deficient animals *glp-1(e2141)* and *glp-4(bn2)*. Experiment conducted at the restrictive temperature 25 °C to deplete the germline. Significance determined by Welch’s unpaired t-test, ****p <0.0001; n = 2. c. Fluorescence comparison of *ucr-2.3(uth252); rgef-1p::Q40::YFP; hsp-6p::GFP* animals on control or *vit-1, vit-3, vit-5* RNAi. Significance determined by Welch’s unpaired t-test, **** p < 0.0001, *** p < 0.001; n > 3.

## Methods

### *C. elegans* strain generation and maintenance

*C. elegans* animals were fed standard OP50 bacteria and grown on standard NGM containing media. For all RNAi knockdown experiments, *C. elegans* animals were fed HT115 bacteria expressing dsRNA targeted to the gene of interest on NGM media containing IPTG and appropriate antibiotics for selection of the dsRNA-containing plasmid. All strains were either generated by this study, generated by SUNY Biotech, or ordered from the Caenorhabditis Genetics Center (CGC). The only exception is the SJZ106 strain (*foxSi27[pie- 1p::tomm20::mKate2::HA::tbb-2 3’UTR]*), which was a kind gift from Dr. Steve Zuryn (The University of Queensland). CRISPR-edited strains were either ordered from SUNY Biotech or generated by this study following standard *C. elegans* CRISPR/Cas9 genome editing protocols (Arribere et al., 2014) and the Alt-R CRISPR-Cas9 system (IDT). Briefly, Cas9 protein (UC Berkeley QB3 MacroLab) was combined with Alt-R CRISPR tracrRNA, a crRNA mixture (targeted to our gene of interest and *dpy-10*), and ssDNA repair template mixture (again, targeted to our gene of interest and *dpy-10*) as a ribonucleoprotein complex and injected into the gonad of L4 stage *C. elegans* hermaphrodites. This allowed us to select “jackpot” broods containing many F1 animals with “roller” or “dumpy” phenotypes to improve efficiency in selection of broods with edited genomes. mosSCI integration of strains were generated following standard procedures (Frøkjær-Jensen et al., 2008). Briefly, constructs of interest were cloned into the pCFJ356 mosSCI plasmid backbone. A DNA mixture of the mosSCI plasmid of interest, the pCFJ601 plasmid (containing the mosSCI transposase), and mCherry expressing plasmids (to eliminate non-integrated, array animals) was injected into the gonad of young, “uncoordinated” EG6703 animals. Resulting progeny were screened for broods that yielded only “mover” animals with no mCherry-expressing arrays.

### Animal synchronization and maintenance

Unless stated, all experiments were performed with synchronization using a standard hypochlorite treatment protocol. Briefly, animals were washed off NGM plates with a M9 solution and bleached with 1.8% sodium hypochlorite solution until all animal carcasses were disintegrated. Resulting eggs were washed five times with M9 solution using centrifugation and eggs were plated on respective plates for the experiments. All experiments were conducted at 20 °C except for germline ablation experiments with temperature sensitive mutants, which were done at the restrictive temperature of 25 °C as indicated in figure legends.

### Mutagenesis screen

*rgef-1p::Q40::YFP; hsp-6p::GFP* animals at the L4 stage were incubated with Ethyl-methyl sulfonate (EMS, Sigma). Five plates of 250 F1 progeny each were selected to screen 2500 genomes. F2 progeny were screened for suppression of *hsp-6p::GFP* signal in the intestine. Identified suppressors were singled onto fresh plates and further screened for homozygous, recessive mutations. Selected suppressor mutants were backcrossed to the parental strain carrying the reporter and polyQ transgenes at least one time.

### Mutation mapping

We followed a SNP-based whole genome sequencing method to identify putative causative mutations in the suppressor mutants (Doitsidou et al., 2010). Briefly, we outcrossed suppressor mutant strains with males from a Hawaiian CB4866 strain containing the *rgef-1p::Q40::YFP* and *hsp-6p::GFP* transgenes. F2 progeny containing the suppression phenotype were selected and allowed to self-fertilize for two generations. Then, genomic DNA was extracted from these suppressor lines using the DNeasy kit (Qiagen) and sent to the Beijing Genomics Institute for WGS on their Illumina HiSeq platform using single 90 nucleotide reads. Genomic data was analyzed using MAQgene as previously described (Bigelow et al., 2009; Doitsidou et al., 2010).

### Fluorescence stereoscope imaging and quantification

Animals were synchronized by hypochlorite treatment and hatched on either control or respective HT115 RNAi bacteria. Animals were grown to Day 2 at 20 °C unless otherwise indicated, like for the restrictive temperature experiments. For imaging experiments in the *rab-3p::hsf-1 OE; hsp- 16.2p::GFP* background, animals were heatshocked for imaging as had been previously published (Gildea et al., 2022). Briefly, animals were hatched and allowed to develop at either 20 °C or 25 °C (if a temperature sensitive mutant). On the day of the experiment, animals were then heatshocked at 34 °C for 2 hr, then let recover at their original temperature they developed at for 2 hr, then imaged. For imaging, D2 animals were anesthesized with 0.1 M sodium azide and lined up on an NGM plate without a bacterial lawn. Animals were imaged using M250FA stereoscope (Leica) under respective fluorescence. Animals were imaged at 250ms exposure under GFP excitation fluorescence unless otherwise indicated. Quantification of intestinal GFP signal was performed using ImageJ/FIJI by tracing the intestinal regions. Statistical analyses were performed with GraphPad PRISM as described in the figure legends.

### Quantitative RT-PCR

Synchronized animals were collected on Day 2 of adulthood and gravity washed in M9 buffer to isolate adults and remove larvae. Animals were collected in three independent biological replicates for one experiment. Washed animals were then frozen in TRIzol (Invitrogen) and freeze thawed using liquid nitrogen. RNA was harvested from animals using a TRIzol-based extraction method and RNA was purified using a RNeasy Mini Kit (Qiagen). cDNA was synthesized using the QuantiTect Reverse Transcription Kit (Qiagen) using equivalent amounts of RNA per sample. qPCR was performed using a standard curve protocol using SYBR Select Master Mix (Life Technologies). Statistical analyses were performed with GraphPad PRISM as described in the figure legends. Primers used are:

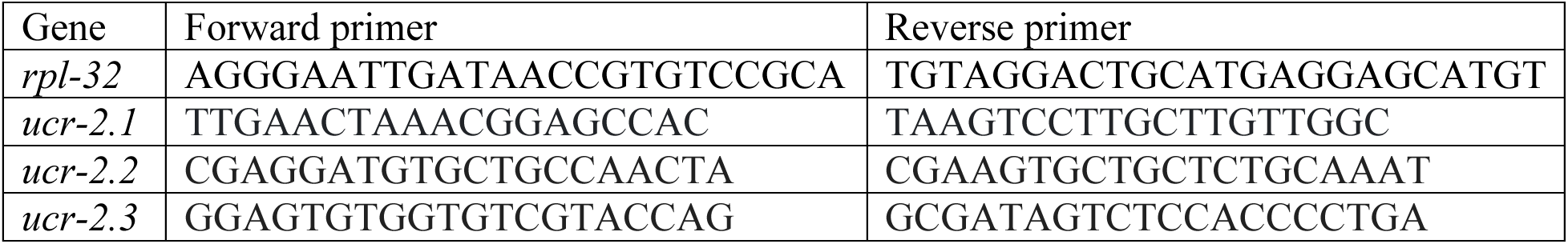

### Sub-cellular fractionation and western blotting

Sub-cellular fractionation and mitochondrial enrichment of *C. elegans* lysates was performed as previously described (Xin et al., 2022). Briefly, synchronized animals were mechanically homogenized with a Dura-Grind Stainless Steel Dounce Trissue Grinder (Wheaton) in mitochondrial extraction buffer (5 mM Tris-HCl / 7.4, 210 mM mannitol, 70 mM sucrose, and 0.1 mM EDTA) with protease inhibitor. Lysates were subject to differential centrifugation. Protein concentration was measured using a Rapid Gold BCA kit (Pierce) before being loaded onto Tris- Glycine 4-12% gels (Invitrogen). Gels were transferred using NuPAGE transfer buffer (Invitrogen) onto nitrocellulose membranes (Biorad) and blocked with LI-COR Blocking Buffer (LI-COR) with Tween. Membranes were probed with antibodies targeted to HA (Abcam, ab9110), NDUFS3 (Abcam, ab14711), and α Tubulin (Sigma, T9026).

### Cell culture, virus generation and transduction

Human hTERT-RPE-1 PAC knockout cells (gifted by Andrew Holland (Lambrus et al., 2016)) were grown in DMEM:F-12 (Gibco) supplemented with 10% FBS (VWR Scientific), 1% Glutamax (Gibco), 1% NEAA (Gibco), and 1% Penicillin/Streptomycin (Gibco). HEK-293T cells were grown in DMEM (GIBCO) containing 10% FBS (VWR Scientific), 1% Glutamax (Gibco), 1% NEAA (Gibco), and 1% Penicillin/Streptomycin (Gibco). Cells were maintained at 37° C with 5% CO2 at atmospheric O2. Cultures were passaged enzymatically every 3-4 days using 0.25% Trypsin (Gibco) diluted 1:3 in 1X PBS (Gibco). Trypsin was inactivated by cell media.

### Cloning of plasmids for cell culture

*C. elegans* (human codon-optimized) UCR-2.3-mNeonGreen and OMP25-tagBFP2 were synthesized as geneblocks (IDT), PCR amplified with Q5 polymerase (New England Biolabs) and incubated with restriction enzyme-digested CD510-B1 plasmid (SystemBio) and assembled with Gibson Assembly Master Mix (New England Biolabs).

### Generation and transduction of UCR-2.3 lentivirus

Lentiviral plasmids encoding pCMV-OMP25-tagBFP2-EF1a-NEO and pUbC-UCR-2.3- mNeonGreen-EF1a-Puro were transfected into HEK-293T cells with lentiviral packaging plasmids with Lipofectamine 3000 (Invitrogen) according to standard lentiviral production protocols. Viral supernatant was filtered through a 0.45 µm filter before transduction. RPE-1 cells were transduced with lentivirus at an MOI of 0.3-0.5 (usually between 50-100 µL of virus/well) and 10 µg/mL Polybrene (Sigma-Aldrich). After 48 hours, RPE-1 cell media was replaced and grown in the presence of 1 µg/mL Puromycin (Gibco) or 100 µg/mL Geneticin (Gibco) for about 7 days and until all non-transduced control cells died.

### Live cell microscopy of human cells

RPE-1 cells were seeded at 1.25 x 10^5^ cells/dish onto Fibronectin-coated plates, 35-mm glass-bottom dishes with #1.5 cover glass (Cellvis) 24 hours prior to imaging. Prior to live cell imaging, cell medium was replaced with pre-warmed FluoroBrite DMEM (Gibco) supplemented with 10% FBS, 1% Glutamax, 1% NEAA, 1% Penicillin/Streptomycin, 1 µM SYTO Deep Red Fluorescent Nuclei Acid Stain (Invitrogen), and 1 µM Probenecid (Invitrogen). Imaging of live cells was performed with a Zeiss Airyscan microscope LSM900 equipped with heated stage and environmental chamber maintained at 37° C and 5% CO2. Images were acquired using the ZenBlue3.1 software followed by deconvolution and Airyscan processing, and further processed using ImageJ/FIJI. Mander’s colocalization coefficients are calculated using the Colocalization threshold plug-in in ImageJ, in which the thresholds were set by the Costes Auto threshold method.

### Mitochondrial oxygen consumption assays

All mitochondrial oxygen consumption rate assays (OCR) were performed using a Seahorse XFe96 Analyzer (Agilent) at 20 °C following a previously published protocol . Briefly, the sensor cartridge was calibrated the day before at room temperature, and appropriate drugs (NaN3, FCCP) were loaded into the injection ports. Day 2 adult animals were gravity washed with M9 buffer, then loaded 10-20 animals per well into the cell culture plate. After respirometry run was completed, the number of animals were counted per well to normalize the data. Statistical analyses were performed with GraphPad PRISM as described in the figure legends.

### Fluorescence widefield microscopy of *C. elegans*

Slides for imaging were prepared by making a fresh flattened 5% agarose pad. Animals were immobilized in 12 μl of 0.1% NaN3, then sealed beneath a 22x22mm coverglass. 6-10 animals were imaged per condition per experiment. Imaging was done using a Leica DM6 Thunder Imager and processed using LAS X software automated Thunder small volume computational clearing.

### Brood size assay

For each strain tested, five L4 animals were singled onto OP50 bacterial plates and kept at 20 °C to lay. Animals were moved each day to new OP50 plates until Day 5. Progeny laid by each singled animal were counted at the L4 or Day 1 life stage. Statistical analyses were performed with GraphPad PRISM as described in the figure legends.

### Quantification of mitochondrial features using FIJI

To quantify mitochondrial number and size in images of mitochondrial reporter worms, Z-stack images were first projected into a single layer using the “Average Intensity” function. The “Auto Local Threshold” function was applied with parameters of “method=Bernsen radius=15 parameter_1=0 parameter_2=0 white” to define mitochondria. The “Skeletonize (2D/3D)” function was applied to extract individual mitochondrion and the “Analyze Skeleton (2D/3D)” function was applied to count the number of these mitochondrial skeletons. To calculate mitochondrial density, the area of the whole germline region was manually selected and measured using FIJI (1 pixel area=0.010645 µm^2^).

### Exogenous serotonin addition

Exogenous serotonin addition was performed following a published procedure (Berendzen et al., 2016). Serotonin hydrochloride (Sigma) was solubilized in MilliQ water and plated onto NGM agar plates already containing control HT115 RNAi bacteria. Solubilized serotonin concentrations were calculated so that the final serotonin concentration as stated considers the total agar volume of the plate (approximately 9.9 ml). Seeded serotonin plates were allowed to dry overnight in the dark and plated with eggs harvested from a hypochlorite treatment for the experiment.

### 5-Fluoro-2’deoxyuridine (FUDR) treatment

Eggs were harvested from hypochlorite treatment of animals and plated on control HT115 bacterial plates. 100 μL of 10 mg/mL FUDR was seeded onto the control HT115 bacterial lawn and let dry overnight at room temperature. At the L4 developmental stage, animals were moved onto the FUDR-treatment plates and let grown to Day 2 of adulthood for their respective experiment.

### Oil Red O and Nile Red staining

Lipid staining using Oil Red O or Nile Red was conducted following previously published protocol (Escorcia et al., 2018). Briefly, synchronized animals were grown to Day 2 on respective HT115 RNAi media. Animals were washed in PBS-T, fixed in 40% isopropanol, then incubated with Oil Red O or Nile Red stain for 2 hours at room temperature shielded from light. Excess dye was washed off and animals were imaged via fluorescence stereoscope imaging for Nile Red using a M250FA stereoscope (Leica) or brightfield/colorimetric imaging for Oil Red O using a Revolve microscope (Echo). Quantification of signal was performed in ImageJ by tracing intestinal regions for fluorescent Nile Red images or Oil Red O colorimetric images. For quantification, Oil Red O images were transformed into the HIS color space using the Color Transformer 2 plugin and quantified in the (S) channel. Statistical analyses were performed with GraphPad PRISM as described in the figure legends.

### RNA-seq library preparation and analysis

RNA was prepared using methods similar to those for RT-qPCR as described above. RNASeq library preparation was performed using Roche products, KAPA Biosystems mRNA HyperPrep Kit and KAPA Unique Dual Index Adapters. Thermo Fisher Scientific NanoDrop and Qubit instruments were used to measure nucleic acid concentrations. Agilent BioAnalyzer was used to determine initial Total RNA and RNASeq library Quality. Single direction sequencing was performed using an Illumina NovaSeq, mode SP, SR100 at the Vincent J. Coates Genomic Sequencing Core at University of California, Berkeley. RNASeq analysis was performed by uploading .fastq files to the Galaxy web platform (The Galaxy Community et al., 2022), using the public server at usegalaxy.org and the *C. elegans* FASTA reference transcriptome Caenorhabditis_elegans.WBcel235.cdna.all.fa, downloaded from ensembl. Tools included Kallisto Quant v0.48.0+galaxy1 and DESeq2, v2.11.40.8+galaxy0(Afgan et al., 2018). GO term enrichment analysis was conducted using the PANTHER Overrepresentation Test on geneontology.org using genes significantly up-regulated with log_2_(FC) > 1. Only results with FDR p value < 0.05 were considered.

